# Common brain networks between major depressive disorder and symptoms of depression that are validated for independent cohorts

**DOI:** 10.1101/2020.04.22.056432

**Authors:** Ayumu Yamashita, Yuki Sakai, Takashi Yamada, Noriaki Yahata, Akira Kunimatsu, Naohiro Okada, Takashi Itahashi, Ryuichiro Hashimoto, Hiroto Mizuta, Naho Ichikawa, Masahiro Takamura, Go Okada, Hirotaka Yamagata, Kenichiro Harada, Koji Matsuo, Saori C Tanaka, Mitsuo Kawato, Kiyoto Kasai, Nobumasa Kato, Hidehiko Takahashi, Yasumasa Okamoto, Okito Yamashita, Hiroshi Imamizu

## Abstract

Many studies have highlighted the difficulty inherent to the clinical application of fundamental neuroscience knowledge based on machine learning techniques. It is difficult to generalize machine learning brain markers to the data acquired from independent imaging sites, mainly due to large site differences in functional magnetic resonance imaging. We address the difficulty of finding a generalizable major depressive disorder (MDD) brain network markers which would distinguish patients from healthy controls (a classifier) or would predict symptom severity (a prediction model) based on resting state functional connectivity patterns. For the discovery dataset with 713 participants from 4 imaging sites, we removed site differences using our recently developed harmonization method and developed a machine learning MDD brain network markers. The classifier achieved 70% generalization accuracy, and the prediction model moderately well predicted symptom severity for an independent validation dataset with 449 participants from 4 different imaging sites. Finally, we found common 2 functional connections between those related to MDD diagnosis and those related to depression symptoms. The successful generalization to the perfectly independent dataset acquired from multiple imaging sites is novel and ensures scientific reproducibility and clinical applicability.

---

A recent initiative, the Research Domain Criteria (RDoC), has sought to redefine and subtype psychiatric disorders in terms of biological systems, without relying on a diagnosis based solely on symptoms and signs. This initiative is expected to inform our understanding of overlapping and heterogeneous clinical presentations of psychotic disorders ^1-4^. In particular, resting state functional magnetic resonance imaging (rs-fMRI) is a useful modality to this end because it enables us to non-invasively investigate whole brain functional connectivity (FC) in diverse patient populations ^5,6^. Rs-fMRI allows for the quantification of the FC of correlated, spontaneous blood-oxygen-level dependent (BOLD) signal fluctuations ^7^. According to the original idea of the RDoC initiative, redefinition and subtyping of psychiatric disorders should be achieved by applying the so-called unsupervised learning technique to the FCs without relying on a diagnosis as a ground truth ^8-10^. However, the number of explanatory variables, FCs, is usually between 10,000 and 100,000, while the sample size, i.e. the number of participants, is usually between 100 and 1,000. Thus, overfitting to noise in the data by machine learning algorithms and the resultant inflation of prediction performance can easily occur unless special precautions are taken ^11^. This situation makes it difficult to directly apply unsupervised learning algorithm to FC data.

To address this problem, we proposed the following hierarchical supervised / unsupervised approach, having partially succeeded in several studies ^12-15^. First, we identified a small number of FCs that reliably distinguish healthy controls (HCs) and psychiatric disorder patients using a supervised learning algorithm. We can use the identified FCs not only for a brain network biomarker of the psychiatric disorder but also for biologically meaningful dimensions of the disorder. Second, we applied unsupervised learning to these low biological dimensions to further understand psychiatric disorders. For instance, we were able to achieve subtyping of major depressive disorder (MDD) by locating MDD patients in these dimensions (subtyping). It may be possible to evaluate the drug effect by locating patients before and after the pharmacological treatment in these biological dimensions ^12^. Furthermore, locating different psychiatric disorder patients (e.g. MDD, schizophrenia [SCZ] and autism spectrum disorder [ASD]) in these dimensions may reveal the relationships among the disorders (multi-disorder spectrum) ^12-15^. In this way, although our approach starts with supervised learning based on diagnosis, our final goal is to understand psychiatric disorders in the biological dimensions while avoiding overfitting to noise in the discovery dataset and ensuring generalization performance for the independent data in completely different multiple imaging sites.

Furthermore, an increasing number of studies have highlighted the difficulty in finding a clear association between existing clinical diagnostic categories and neurobiological abnormalities ^8,16,17^. Therefore, the necessity of a symptom-based approach, which directly describes the association with neurobiological abnormalities, is increasingly recognized rather than a diagnosis-based approach ^18^. Here, we also construct a brain network marker which would predict symptom severity.

Whether a brain network marker constructed in the first stage generalizes to the data acquired from multiple completely different imaging sites is a very important issue for the above hierarchical supervised/unsupervised approach ^19-21^. However, an increasing number of studies have highlighted the difficulty in generalization of the brain network marker to the data acquired from multiple completely independent imaging sites, even using the supervised learning method ^22,23^. For example, in a recent paper by Drysdale, which is one of the most successful brain network markers of MDD, the classification accuracy for MDD in completely independent imaging sites was 68.8% for 16 patients from 1 site, which represents only 3% of the validation cohort (Drysdale et al., 2017, Supplementary Tables 3 and 6).

Here, we targeted MDD, the world’s most serious psychiatric disorder in terms of its social repercussions ^24,25^, and investigated whether we could construct a brain network marker which generalizes to the data acquired from multiple completely different imaging sites. We considered and satisfied 3 issues and conditions to ensure generalization of our network marker of MDD to the independent validation dataset, which does not include imaging sites of the discovery dataset. First, we used our recently developed harmonization method, which could reduce site differences in FC ^26^. According to our recent study, the differences in resting state FCs for different imaging sites consist of measurement bias due to differences in fMRI protocols and MR scanners, and sampling bias due to recruitment of different participant populations. The magnitude of the measurement bias was larger than the effects of disorders including MDD, and the magnitude of the sampling bias was comparable to the effects of disorders ^26^. Therefore, a reduction in the site difference in FC is essential for the generalization of network models in the validation dataset. Second, we validated our network marker using a perfectly independent and large cohort collected from multiple completely different imaging sites from a Japanese nation-wide database project called the Strategic Research Program for Brain Science (https://bicr.atr.jp/decnefpro/). We used a rs-fMRI discovery dataset with 713 participants (149 MDD patients) from 4 imaging sites and an independent validation dataset with 449 participants (185 MDD patients) from 4 imaging sites that were not included in the discovery dataset. We further used another dataset of 75 HCs, 154 SCZ patients and 121 ASD patients to investigate the multi-disorder spectrum. In total, we used 1,512 participants’ data in this study. Furthermore, unlike previous studies that restricted the subtype of MDD ^9,12^, we targeted all MDD patients without restricting according to depression subtype in order to enable future subtyping in the biological dimensions, which is beyond the purpose of the current paper. Third, we carefully avoided overfitting noise in the discovery dataset. As explained above, the number of explanatory variables is typically larger than the sample size in the rs-fMRI study, thus overfitting to noise in the discovery dataset by machine learning algorithms and resultant inflation of prediction performance can happen easily unless special precautions are taken. We used a sparse machine learning algorithm with the least absolute shrinkage and selection operator (LASSO) to avoid overfitting to noise and selected only essential FCs ^27^. As a result, for the first time, to our knowledge, we developed a generalizable brain network marker for MDD diagnosis without restricting to certain subtypes such as treatment-resistant or melancholy types and a generalizable brain network marker for depression symptoms.

## Results

### Datasets

We used two rs-fMRI datasets for the analyses. The “discovery dataset contained data from 713 participants (564 HCs from 4 sites, 149 MDD patients from 3 sites; Table 1), and the “independent validation dataset” contained data from 449 participants (264 HCs from independent 4 sites, 185 MDD patients from independent 4 sites; Table 1). Most data utilized in this study can be downloaded publicly from the DecNef Project Brain Data Repository (https://bicr-resource.atr.jp/srpbsopen/ and https://bicr.atr.jp/dcn/en/download/harmonization/). The imaging protocols and data availability statement of each site is described in Supplementary Table 1. Depression symptoms were evaluated using the Beck Depression Inventory-II (BDI-II) score obtained from most participants in each dataset. Clinical details such as medication information and the presence of comorbidities in patients with MDD are described in Supplementary Table 2.

**Table 1.**
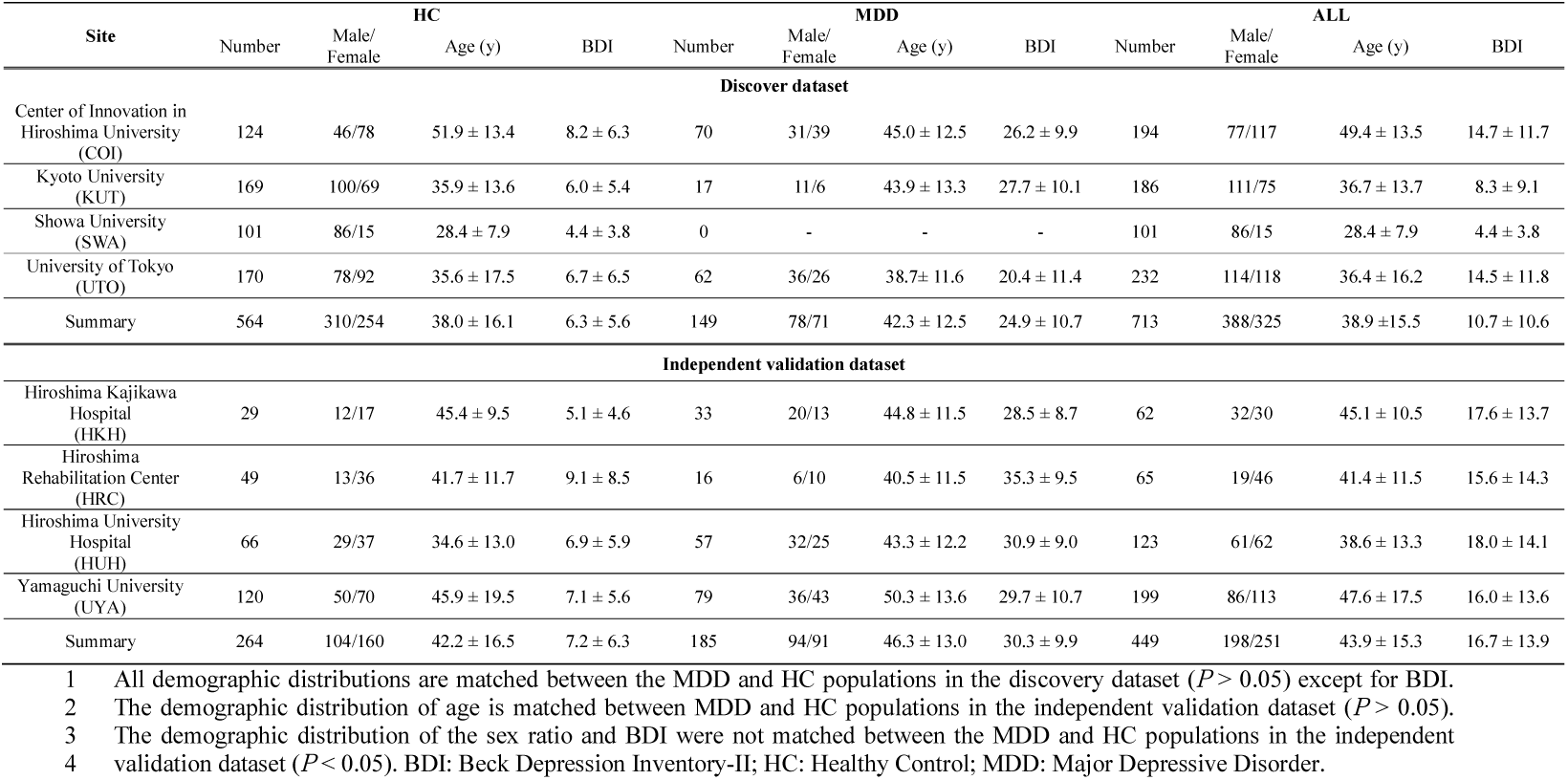
Demographic characteristics of participants in both datasets.

### Site difference control in FC

Classical preprocessing was performed, and FC was defined based on a functional brain atlas consisting of 379 nodes (regions) covering the whole brain ^28^. The Fisher’s *z*-transformed Pearson correlation coefficients between the preprocessed BOLD signal time courses of each possible pair of nodes were calculated and used to construct 379 x 379 symmetrical connectivity matrices in which each element represents a connection strength, or edge, between two nodes. We used 71,631 connectivity values (379 x 378 / 2) of the lower triangular matrix of the connectivity matrix. To control for site differences in the FC, we applied a traveling subject harmonization method to the discovery dataset ^26^. In this method, the measurement bias (the influence of the difference in the properties of MRI scanners, such as the imaging parameters, field strength, MRI manufacturer, and scanner model) was estimated by fitting the regression model to the FC values of all participants from the discovery dataset and the traveling subject dataset, wherein multiple participants travel to multiple sites to assess measurement bias (see *Control of site differences* in Methods section). This method enabled us to subtract only the measurement bias while leaving important information due to differences in subjects among imaging sites. We applied the ComBat harmonization method ^29-32^ to control for site differences in the FC of the independent validation dataset because we did not have a traveling subject dataset for those sites.

### Reproducible FCs related to MDD diagnosis and depression symptoms

Utilizing a simple mass univariate analysis, we investigated the reproducibility of the effect sizes by diagnosis and depression symptoms on individual FC across the discovery and validation datasets. For the effect of the diagnosis on each FC, we calculated the difference in the FC value across participants between the HCs and the MDDs (*t*-value). The Pearson’s correlation coefficient between FC strength and BDI scores (*r*-value) was calculated for the effect of the depressed symptom on each FC. Fig. 1a left scatter plot shows the diagnosis effect size for the discovery dataset in the abscissa and that for the validation dataset in the ordinate for each FC. Fig. 1 a right is a scatter plot for the symptom effect size. Effect sizes for the two datasets were positively correlated, implying reproducibility of these effects. We compared the distributions of diagnosis and symptom statistics of the discovery dataset to the distributions in the shuffled data in which diagnosis and symptom severity were permuted across subjects. We found the larger effects of the diagnosis in the original data in comparison to the shuffled data (Fig. 1a left histograms). We confirmed that the results were similar for the symptom (Fig. 1a right histograms). These results indicate that resting-state FCs contain consistent information across the two datasets regarding MDD diagnosis and depression symptoms.

**Figure 1:**
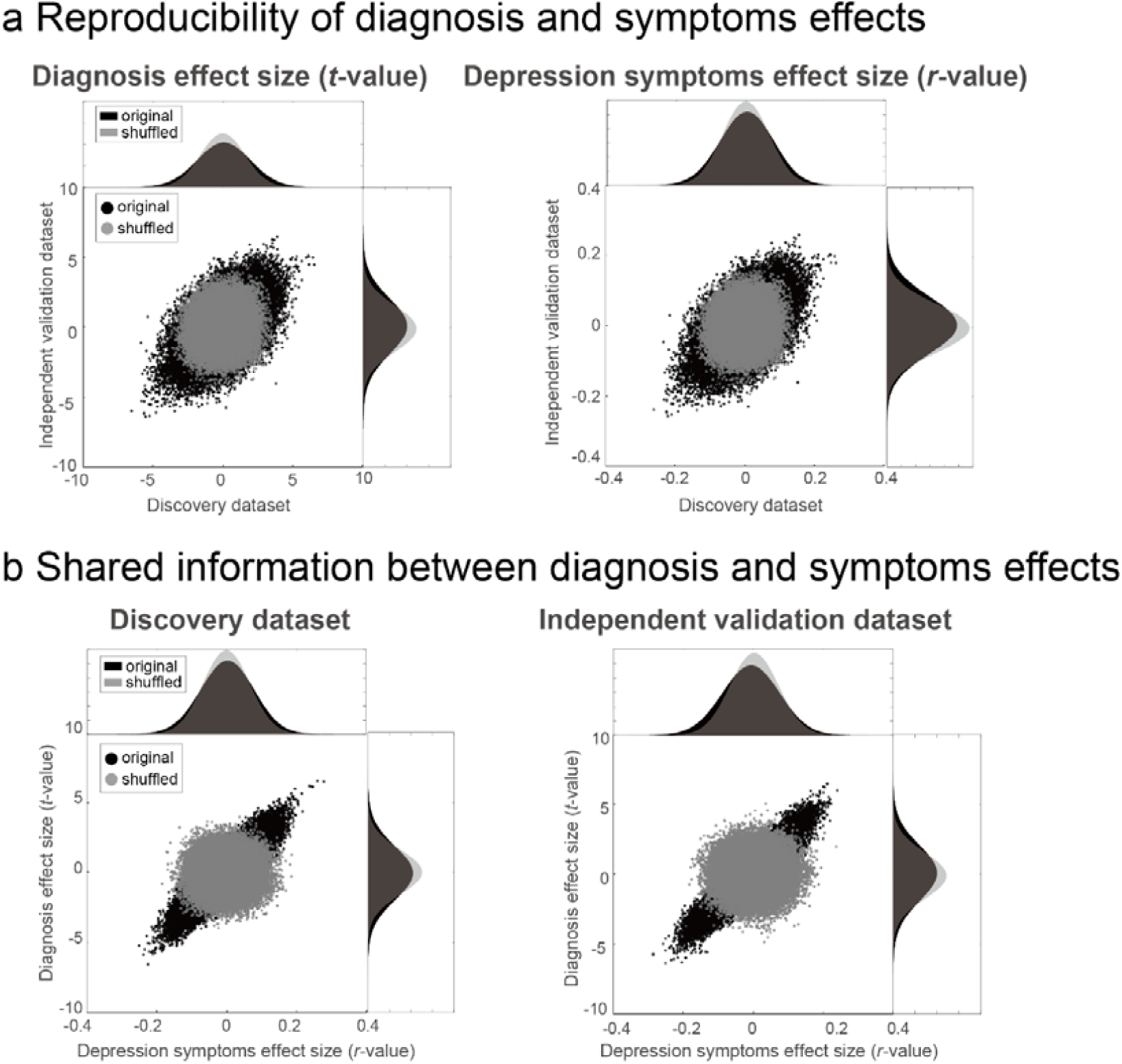
Results of mass univariate analysis. (a) Reproducibility across the two datasets regarding diagnosis (left) and symptom (right) effects. (left) Scatter plot and its histograms of the diagnosis effect size (the difference in mean functional connectivity strengths between depressed patients and healthy groups: *t*-value). Each point in the scatter plot represents the diagnosis effect in the discovery dataset in the abscissa and that for the independent validation dataset in the ordinate for each functional connectivity. (right) Same format for the depression symptoms effect size (Pearson’s correlation between BDI-II and functional connectivity strength: *r*-value). The original data is in black, while the shuffled data in which subject information was permuted is in gray. (b) Shared information between diagnosis and symptom effects. Scatter plots and its histogram of the diagnosis effect size (*t*-value) in the ordinate and the depression symptoms effect size (*r*-value) in the abscissa for all functional connectivity within the discovery dataset (left) and the validation dataset (right). Each point represents symptom effect size in the abscissa and that for diagnosis in the ordinate for each functional connectivity. The original data is in black, while the shuffled data in which subject information was permuted is in gray.

Furthermore, to statistically evaluate the reproducibility of these effects on FCs, we calculated Pearson’s correlation between the discovery and validation datasets regarding the above two statistics (*t*-values for diagnosis and *r*-values for symptom). We found significant correlations between the two datasets for diagnosis (*t*-value: *r*_(71631)_ = 0.51, 95 % confidence interval (CI) = [0.508 0.519], *R*^*2*^*=*0.26, [permutation test, *P* < 0.001, one-sided]), as well as for symptom (*r*-value: *r*_(71631)_ = 0.39, 95 % CI = [0.380 0.393], *R*^*2*^*=*0.15, [permutation test, *P* < 0.001, one-sided], Fig. 1a). This result indicates that the effects of MDD diagnosis on FCs and the effects of symptom severity were reproducible even in the independent dataset acquired from completely different sites.

### Shared information on FCs between MDD diagnosis and depression symptoms

We investigated whether the FCs related to MDD diagnosis and the FCs related to depression symptoms share identical information or partially overlapping information. To this end, we calculated Pearson’s correlation between the *t*-values and *r*-values on FCs in the same dataset. We found high correlations but not completely identical (Discovery dataset: *r* = 0.86, Independent validation dataset: *r* = 0.91, Fig. 1b). This result indicates that shared information exists on FCs between MDD diagnosis and depression symptoms.

### Brain network marker of MDD diagnosis generalized to MDD data obtained from completely different multisites

We constructed a brain network marker for MDD, which distinguished between HCs and MDD patients, using the discovery dataset based on 71,631 FC values. Based on our previous studies ^12-15,33^, we assumed that psychiatric disorder factors were not associated with whole brain connectivity, but rather with a specific subset of connections. Therefore, we used logistic regression with LASSO, a sparse machine learning algorithm, to select the optimal subset of FCs ^34^. We have already succeeded in constructing generalizable brain network markers of ASD, melancholic MDD, SCZ and obsessive compulsive disorder ^12-15,33^ by using a similar sparse estimation method that automatically selects the most important connections. We also tried a support vector machine (SVM) for classification instead of LASSO. However, the performance was not improved compared to that with LASSO (Supplementary Note 1).

To estimate the weights of logistic regression and a hyperparameter that determines how many FCs were used, we conducted a nested cross validation procedure (Fig. 2) (see *Constructing MDD classifier using the discovery dataset* in the Methods section). We first divided the whole discovery dataset into the training set (9 folds out of 10 folds), which was used for training a model, and the test set (1 fold out of 10 folds), which was used for testing the model. To avoid bias due to the difference in the numbers of MDD patients and HCs, we used an undersampling method for equalizing the numbers between the MDD and HC groups ^35^. Since only a subset of training data is used after undersampling, we repeated the random sampling procedure 10 times (i.e., subsampling). We then fitted a model to each subsample while tuning a regularization parameter in an inner loop of nested cross validation, resulting in 10 classifiers. The mean classifier-output value (diagnostic probability) was considered indicative of the classifier output. Diagnostic probability values > 0.5 were considered to be indicative of a MDD diagnosis. We calculated the area under the curve (AUC), accuracy, sensitivity, specificity, positive predictive value (PPV), and negative predictive value (NPV). Furthermore, we evaluated classifier performance for the unbalanced dataset using the Matthews correlation coefficient (MCC) ^36,37^, which takes into account the ratio of the confusion matrix size.

**Figure 2:**
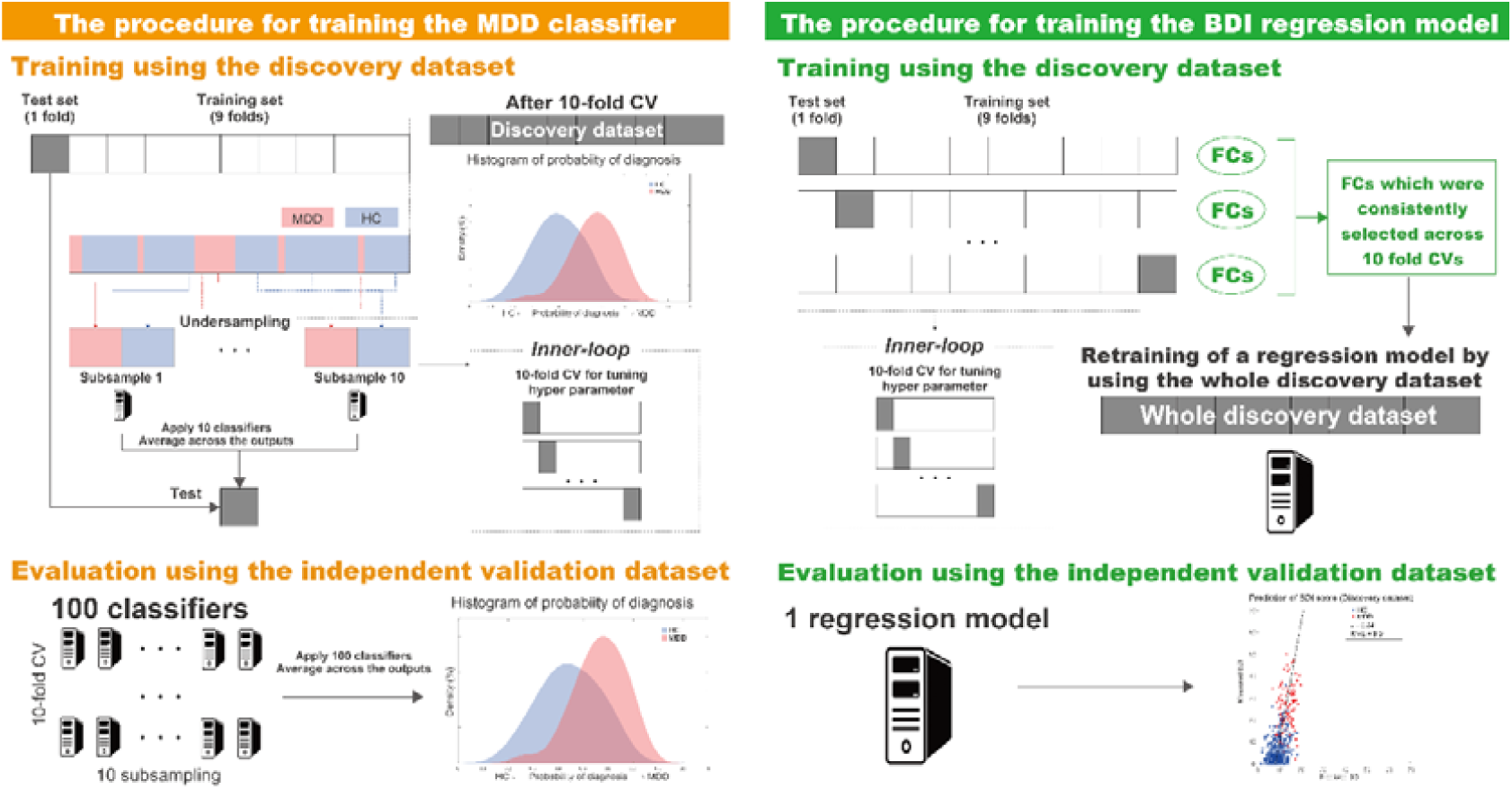
Schematic representation of the procedure for training brain network markers and evaluation of their predictive power. The MDD classifier was constructed using a nested cross validation procedure, undersampling, and subsampling technique in the discovery dataset. The BDI regression model was constructed using the union of FC values selected by the embedded method in the discovery dataset. Generalization performances were evaluated by applying the constructed classifiers and to the independent validation dataset. The machine learning classifiers are represented by PC cartoons. BDI: Beck Depression Inventory-II, CV: cross validation, MDD: major depressive disorder, HC: healthy control, FC: functional connectivity.

The classifier distinguished MDD and HC populations with an accuracy of 67% in the discovery dataset. The corresponding AUC was 0.77, indicating acceptable discriminatory ability. Fig. 3a shows that the two diagnostic probability distributions of the MDD and HC populations were clearly separated by the 0.5 threshold (right, MDD; left, HC) for the discovery dataset. The sensitivity, specificity, PPV, and NPV were 75%, 65%, 0.35, and 0.91, respectively. This classifier led to an acceptable MCC of 0.33. We found that acceptable classification accuracy was achieved for the full dataset as well as for the individual datasets from 3 of the imaging sites (Fig. 3b) to similar degrees. Only HC individuals were identified in the SWA dataset; however, notably, its probability distribution was comparable to the HC populations at other sites.

**Figure 3:**
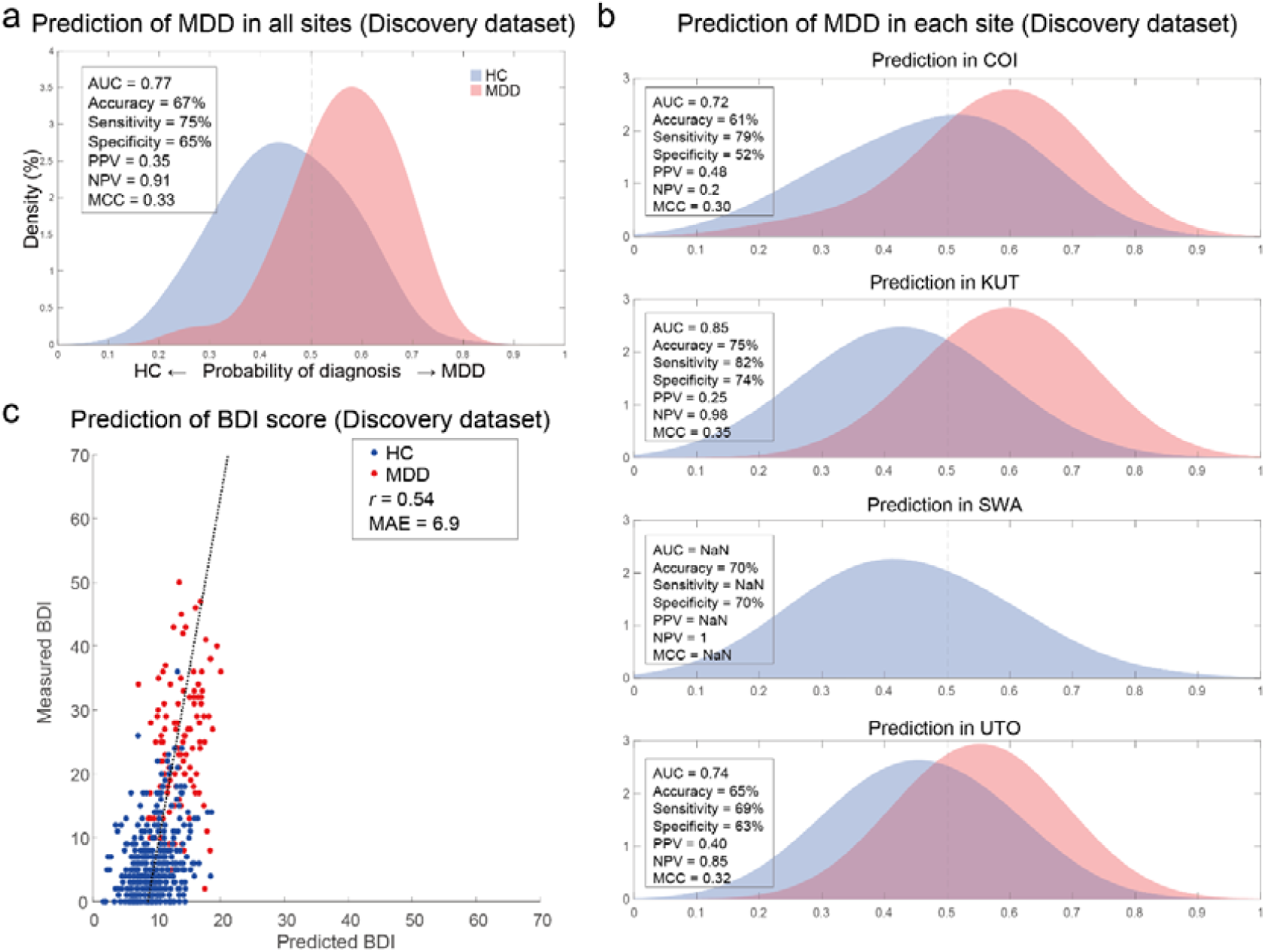
MDD classifier and BDI regression model performances in the discovery dataset. (a) The probability distribution for the diagnosis of MDD in the discovery dataset and (b) probability distributions for each imaging site. MDD and HC distributions are depicted in red and blue, respectively. (c) Scatter plots of measured and predicted BDI. The solid line indicates the linear regression of the measured BDI from the predicted BDI. The correlation coefficient (*r*) and mean absolute error (MAE) are shown. Each data point represents one participant. BDI: Beck Depression Inventory-II; HC: healthy control; MDD: major depressive disorder; AUC: area under the curve; PPV: positive predictive value; NPV: negative predictive value; MCC: Matthews correlation coefficient; COI: Center of Innovation in Hiroshima University; KUT: Kyoto University; SWA: Showa University; UTO: University of Tokyo.

We tested the generalizability of the classifier using an independent validation dataset. We created 100 classifiers of MDD (10-fold × 10 subsamples); therefore, we applied all trained classifiers to the independent validation dataset. Next, we averaged the 100 outputs (diagnostic probability) for each participant and considered the participant as a patient with MDD if the averaged diagnostic probability value was > 0.5. The classifier distinguished the MDD and HC populations with an accuracy of 69% in the independent validation dataset. If the accuracy for the validation dataset is much smaller than that of the discovery dataset, overfitting is strongly suggested and the reproducibility of the results is put into doubt. In our case, 69% accuracy for the validation dataset was actually higher than 67% accuracy for the discovery dataset, and this concern does not apply. The corresponding AUC was 0.77 (permutation test, *P <* 0.01, one-sided), indicating an acceptable discriminatory ability. Fig. 4a shows that the two diagnostic probability distributions of the MDD and HC populations were clearly separated by the 0.5 threshold (right, MDD; left, HC). The sensitivity, specificity, PPV, and NPV were 74%, 65%, 0.60, and 0.78, respectively. This approach led to an acceptable MCC of 0.38 (permutation test, *P <* 0.01, one-sided). In addition, acceptable classification accuracy was achieved for the individual datasets of the 4 imaging sites (Fig. 4b). To investigate whether our classifier can be generalized to milder depression, we applied our classifier to MDD patients with low BDI scores (score <= 20, n = 30) in the independent validation dataset. As a result, 22 of the 30 patients were correctly classified as having MDD (accuracy of 73%), a similar performance level to when the classifier was applied to all patients with MDD. Furthermore, all patients with MDD at the KUT imaging site were treatment-resistant patients (treatment-resistant depression: adequate use of two or more antidepressants for 4-6 weeks is not efficacious, or intolerance to two or more antidepressants exists). We calculated the classification accuracy only at KUT and obtained the same performance level (accuracy = 75%). These results suggest that the current MDD classifier can be generalized to milder depression, as well as to treatment-resistant patients with MDD.

**Figure 4:**
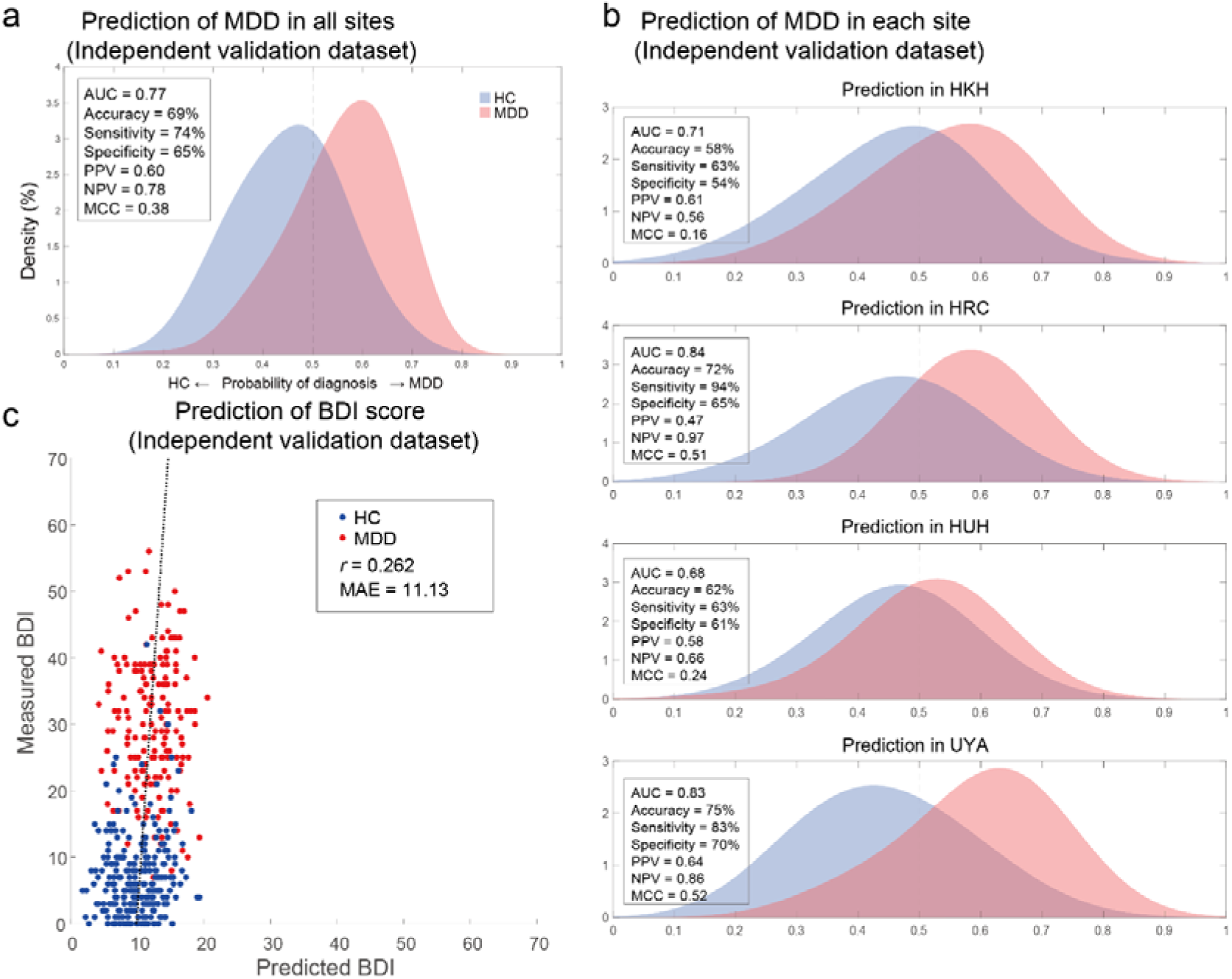
MDD classifier and BDI regression model performances in the independent validation dataset. (a) The probability distribution for MDD diagnosis in the independent validation dataset and (b) probability distributions for each imaging site. MDD and HC distributions are depicted in red and blue, respectively. (c) Scatter plots of measured and predicted BDI. The solid line indicates the linear regression of the measured BDI from the predicted BDI. The correlation coefficient (*r*) and mean absolute error (MAE) are shown. Each data point represents one participant. BDI: Beck Depression Inventory-II; HC: healthy control; MDD: major depressive disorder; AUC: area under the curve; PPV: positive predictive value; NPV: negative predictive value; MCC: Matthews correlation coefficient; HKH: Hiroshima Kajikawa Hospital; HRC: Hiroshima Rehabilitation Center; HUH: Hiroshima University Hospital; UYA: Yamaguchi University.

Regarding the effectiveness of the developed network marker, although discriminability was acceptable (AUC = 0.77) in the independent validation dataset, the performance of the PPV was low in the discovery dataset (0.35). This occurred because the number of patients with MDD was much smaller than that of HCs (about 4 times as many HCs as MDDs) in the discovery dataset. By contrast, in the independent validation dataset, in which the number of HCs is about 1.5 times as high as the number of MDDs, the PPV, at 0.60, was acceptable. When applying a developed network marker in clinical practice, we assume this marker to be applied to those who actually visit the hospital. Therefore, the actual PPV will be acceptable in clinical practice because the prevalence of MDD may be relatively high compared to the general prevalence of MDD. Furthermore, in the independent validation dataset, when we divided the dataset into low- and high-risk groups based on the cutoff value (probability of MDD being 0.51) determined in the discovery dataset ^38^, the odds (sensitivity / 1-sensitivity) were 1.58 in the high-risk group. Moreover, the odds ratio was 5.8 when the odds in the low group were set to 1. That is, the output of the classifier (probability of MDD) will be useful information for psychiatrists as a physical measure supplementing the patients’ symptoms and signs in order to make a diagnosis.

We further investigated whether the discrimination performances were different across imaging sites in the independent validation dataset. We calculated the 95% confidence intervals (CIs) of the discrimination performances (AUC, accuracy, sensitivity, and specificity) using a bootstrap method for every imaging site. We repeated the bootstrap procedure 1,000 times and calculated the 95% CI for each site. We then checked whether there was a site whose CI did not overlap with the CIs of other imaging sites. We were unable to find such an imaging site, suggesting no significant systematic difference. However, we noted that the sensitivity at the HUH site was inferior to that at the two other imaging sites (see Supplementary Note 2 and Supplementary Fig. 1: CI of sensitivity in the HUH does not overlap with CI in the HRC or UYA). We discuss the differences in performances among imaging sites in the Discussion section.

Finally, we checked the stability of our developed network marker to see if the same subject was consistently classified in the same class when the subject was scanned multiple times at various imaging sites. We applied our marker to a traveling subject dataset (Supplementary Table 6) in which 9 healthy participants (all male participants; age range, 24–32 years; mean age, 27 ± 2.6 years) were scanned about 50 times at 12 different sites, producing a total of 411 scan sessions. We achieved a high accuracy in this dataset (mean accuracy = 84.5, 1SD = 12.8, across participants). This result indicates that our developed network marker has high stability even if the same subject is scanned multiple times at various imaging sites.

### Brain network prediction model of depression symptoms generalized to completely different multisite data

We constructed a brain network prediction model of the BDI score using the discovery dataset based on 71,631 FC values. We employed linear regression using the LASSO method. At first, we tried to evaluate the prediction accuracy on the discovery dataset using a 10-fold CV procedure following our method for the MDD classifier. However, no FC was selected by the LASSO in 7 out of 10 folds during the hyperparameter determination. This result indicates that the regularization in the LASSO worked too strongly. Therefore, we constructed a regression model using the FC values selected by the embedded method in the discovery dataset (Fig. 2) ^39^. This approach caused information leakage because we evaluated the model using the discovery dataset, 30% of which was used for selecting the FCs; therefore, the results in the discovery dataset may be overfitted. This reservation meant that it was important to confirm generalization performance by applying this regression model to an independent validation dataset, as described below. Finally, we calculated the mean absolute error (MAE) and Pearson’s correlation coefficients between the predicted and measured BDI scores. The BDI score was well predicted with a significant correlation (*r*_(477)_ = 0.54, 95 % CI = [0.473 0.601], *R*^*2*^*=*0.29, *P* = 1.6 × 10^−37^, one-sided; MAE = 6.9; Fig. 3c). Furthermore, a significant correlation was achieved for HC and MDD populations separately (HC, *r*_(367)_ = 0.28, 95 % CI = [0.185 0.374], *R*^*2*^*=*0.08, *P* = 3.7 × 10^−8^, one-sided; MDD, *r*_(110)_ = 0.30, 95 % CI = [0.116 0.459], *R*^*2*^*=*0.09, *P* = 0.0016). Once again, cautiously, these results may be overfitted because the evaluation data are not independent data. The correct assessment should be based on results from the following independent validation dataset.

We tested the generalizability of the regression model using the independent validation dataset. We created one BDI regression model using all the discovery dataset samples; therefore, we applied the trained regression model to the independent validation dataset and considered its output as the predicted BDI score. The BDI score was moderately well predicted, with a significant correlation in the independent validation dataset (*r* = 0.26; MAE = 11.1; Fig. 4c; permutation test, *P* < 0.01, one-sided). We could not construct any regression model for the whole permutation procedure because no FC were selected at the nested CV in the LASSO procedure. This result indicated that the performance of the BDI regression model in the independent validation data without permutation was statistically significant.

### Important FCs for the brain network markers

We examined important resting state FCs for MDD diagnosis and depression symptoms by extracting the important FCs related to the MDD classifier and BDI regression model, respectively. Briefly, we counted the number of times an FC was selected by LASSO during the 10-fold CV. We considered this FC to be important if this number was significantly higher than the threshold for randomness, according to a permutation test. For the MDD classifier, we permuted the diagnostic labels of the discovery dataset and conducted a 10-fold CV and 10-subsampling procedure, and we repeated this permutation procedure 100 times. We then used the number of counts for each connection selected by the sparse algorithm during 10 subsamplings x 10-fold CV (max 100 times) as a statistic in every permutation dataset. To control for the multiple comparison problem, we set a null distribution as the max distribution of the number of counts over all FCs and set our statistical significance to a certain threshold (permutation test, *P* < 0.05, one-sided). We also performed a permutation test for the BDI regression model. We permuted the BDI scores of the discovery dataset, conducted a 10-fold CV, and repeated this permutation procedure 100 times.

Figures 5a and 5b shows the spatial distribution of the 31 FCs and 13 FCs related to the MDD diagnosis and depression symptoms, respectively, that were automatically and unbiasedly identified from the data by the machine learning algorithms. Two FCs were common between the diagnosis and symptom models. These connections were the connection ① between right insula and right frontal medial orbital cortex, and ② between the right insula and right cingulum anterior cortex. A detailed list of the FCs is provided in Supplementary Tables 3 and 4. We discussed details of these FCs in the Discussion section.

**Figure 5:**
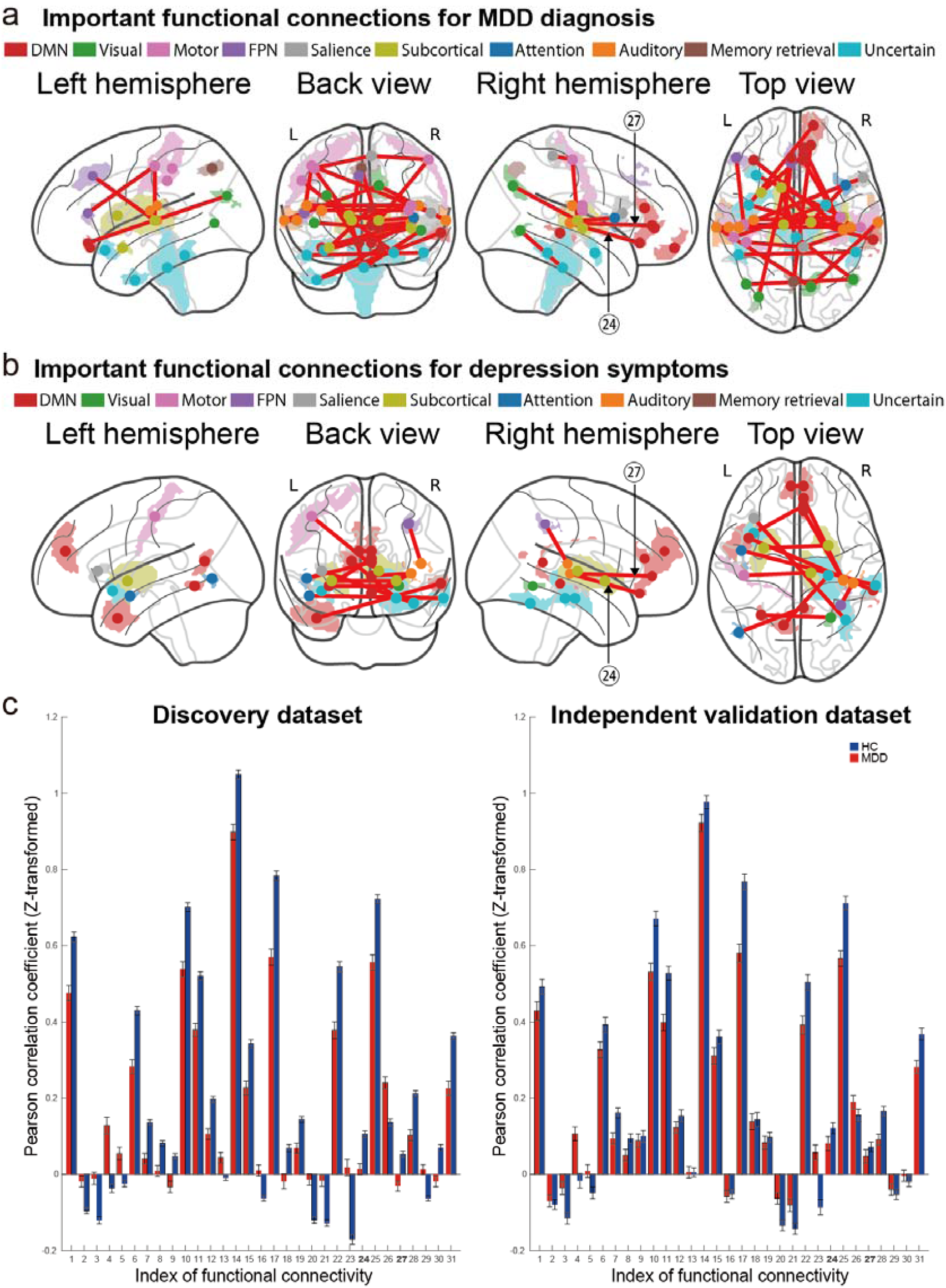
Important FCs for MDD diagnosis and depression symptoms. **(a)** The 31 functional connections (FCs) which are important for MDD diagnosis viewed from left, back, right, and top. Interhemispheric connections are shown in the back and top views only. Regions are color-coded according to the intrinsic network. **(b)** The 13 FCs which are important for depression symptoms viewed from left, back, right, and top. Interhemispheric connections are shown in the back and top views only. Regions are color-coded according to the intrinsic network. Two connections were common (▪ between the right insula and the right frontal medial orbital cortex, and ▪ between the right insula and the right cingulum anterior cortex). **(c)** The FC values of 31 FCs for both HCs and MDD patients in the discovery dataset and the independent validation dataset. MDD: major depressive disorder; DMN: default mode network; FPN: fronto-parietal network.

## Discussion

In the present study, we thoroughly considered conditions and resolved difficulties in order to ensure the generalization of our brain network marker in the independent validation dataset, which does not include any imaging sites of the discovery dataset. We succeeded in generalizing our network marker to the big independent validation dataset. This generalization ensures scientific reproducibility and the clinical applicability of rs-fMRI. Without this fundamental evidence, we cannot proceed to the development of rs-fMRI-based subtyping, evaluation of drug effects, or exploration of multi-spectrum disorder in the biological dimensions, as mentioned in the Introduction section. Therefore, our study found generalizable psychiatric biomarkers which the fields of psychiatry, neuroscience and computational theory have long sought out, to no avail, since the RDoC initiative.

We developed generalizable brain network markers without restriction to treatment-resistant or melancholy MDD types. Most previous studies have reported the performance of a prediction model using data from the same imaging sites using a CV technique. However, because of large imaging-site differences in rs-fMRI data ^26,40^, CV methods generally induce inflations in performance. To ensure reproducibility, it is critical to demonstrate the generalizability of the models with an independent validation dataset acquired from completely different imaging sites ^11,19-21^. To overcome the above-mentioned site differences, we reduced site differences in a multisite large-scale rs-fMRI dataset using our novel harmonization method. Next, we constructed an MDD classifier that was acceptably generalized to the independent validation dataset. Acceptable generalized prediction performance was also achieved for the 4 individual imaging site datasets (Fig. 4b). This generalization was achieved even though the imaging protocols in the independent validation datasets were different from the discovery dataset. There are only two studies in which generalization of FC-based MDD classifiers to independent validation data was demonstrated ^9,12^. To the best of our knowledge, our work is the first to construct a generalized classifier of MDD without restriction to certain MDD subtypes: Drysdale concentrated MDD patients who were treatment resistant and Ichikawa restricted patients with the melancholic subtype of MDD. Constructing the whole MDD marker is important for subsequent MDD subtyping analyses. This was achieved for the first time by collecting data on a large variety of MDD patients from multiple imaging sites and objectively harmonizing them with a traveling subject dataset.

With respect to site differences in prediction performance, we found that the sensitivity at the HUH site was inferior to that of the two other imaging sites (see Supplementary Fig. 1). The reason for which the sensitivity was low at the HUH site is that the threshold for separating HCs and patients with MDD was shifted to the MDD side (the probability of diagnosis = 0.5: see the vertical line in Fig. 4b). In contrast, the thresholds were shifted to the HC side at the HKH and UYA sites. These shifts may be due to the fact that the removal of site differences was insufficient. If FC includes a measurement bias, which represents the site difference, the threshold will shift. Our previous study showed that the measurement bias, which includes the site difference, was the largest at the HUH site ^26^. These results indicate that it is important to remove site differences using precise harmonization methods, such as the traveling subject harmonization method if possible, when we apply a classifier to new subjects collected from a new imaging site. Because of the absence of the traveling subject dataset, traveling subject harmonization was not possible for the independent validation dataset, and we were forced to use ComBat in this case.

The machine learning algorithms reliably identified the 31 FCs which are important for MDD diagnosis (Fig. 5a, and Supplementary Table 3). We hereafter summarize the characteristics of the 31 FCs. First, 25 of 31 FCs exhibited hypo-connectivity in the MDD population in the independent validation dataset (the absolute value of the FC was closer to 0 in MDD than in HC individuals; Fig. 5c). The connectivity between the left and right insula had the largest difference among 31 FCs between MDD patients and HCs (FC 17 in Fig. 5c). Abnormalities in the insula were found not only in MDD patients ^41,42^ but also reported as common abnormalities (reduced gray-matter volume) among psychiatric disorders ^1^. Therefore, the connectivity associated with the insula is a potential candidate for the neurobiological dimension to understand a multi-spectrum disorder. Second, only 3 FCs (FCs 13, 16, and 26) exhibited hyper-connectivity in the MDD population (the absolute value of the FC was greater in MDD than HC individuals) in the independent validation dataset. However, the differences in those 3 FC values between HC and MDD were not significant (Supplementary Table 3). Finally, only 3 FCs (FCs 4, 5, and 23 in Fig. 5c) had reversed FC values between MDD patients and HCs (positive values in MDD patients and negative values in HCs). Two of 3 FCs were the FCs between the right postcentral cortex and the right thalamus, and the left postcentral cortex and the left thalamus. In the study of Drysdale et al.^9^, the FCs related to the thalamus showed stronger and more positive connectivity in MDD patients as a common feature across the 4 biotypes of MDD, consistent with our current results. A previous study also reported increased FC values between the thalamus and sensory motor cortex (postcentral cortex) in MDD patients ^43^. Participants who have an increased FC value between the thalamus and sensory motor cortex have a greater decline in cognitive function and affective experience ^43^. Finally, two FCs were common between the diagnosis and symptom models. These connections were the connection between right insula and right frontal medial orbital cortex, and between the right insula and right cingulum anterior cortex. We need further analyses to clarify how abnormalities in each FC are associated with cognitive and affective functions in a future study.

Ultimately, it would be very important to understand the relationships across disorders (multi-disorder spectrum). For example, investigating the heterogeneous clinical presentations of psychiatric disorders, the MDD-ness, which is the output of the MDD classifier, may provide a useful biological dimension across the multiple-disorder spectrum. To explore this possibility, we applied our MDD classifier to SCZ patients and ASD patients included in the DecNef Project Brain Data Repository (https://bicr-resource.atr.jp/srpbsopen/). We found that SCZ had a high tendency (similarity) toward MDD while ASD had no such a tendency toward MDD (Supplementary Figure 2a). This result suggests that the MDD classifier generalizes to SCZ but not to ASD. We note that our discovery dataset for the construction of the MDD classifier included no patients with MDD who were comorbid with SCZ and only 1 patient with MDD who was comorbid with ASD. Therefore, our classifier was not affected by either SCZ or ASD diagnosis. Thus, the above generalization of the MDD classifier may point to a certain neurobiological relevance among diseases. Our SCZ patients were in the chronic phase and had negative symptoms. Considering that the negative symptoms of schizophrenia are similar to depression symptoms, the generalization hypothesizes the existence of neurobiological dimensions underlying the common symptoms between SCZ and MDD. For example, anhedonia exists as a transdiagnostic symptom between SCZ and MDD ^44,45^, and some studies have been conducted to understand the neurological basis of anhedonia across psychiatric disorders including SCZ and MDD ^45,46^. We need further analyses to quantitatively examine this hypothesis and investigate the neurobiological relationship between SCZ and MDD by gathering more precise information on SCZ (symptoms and medication history). To further understand the multi-disorder spectrum, we developed markers of SCZ and ASD using the same method as in this study in addition to a brain network marker of MDD (Supplementary Note 3). As a result, we found an interesting asymmetric relationship among these disorders: the classifier of SCZ did not generalize to patients with MDD (Supplementary Figures 2b). This kind of asymmetry in the classifiers had also been found between the SCZ classifier and the ASD classifier (the ASD classifier generalized to SCZ, but the SCZ classifier did not generalize to ASD) ^13-15^. These results provide us with important information for understanding the biological relationships between diseases. For example, the above asymmetry between the SCZ and ASD or MDD classifiers suggests that the brain network related to SCZ is characterized by a larger diversity than that of ASD or MDD, and that it partially shares information with the smaller brain network related to ASD or MDD than that of SCZ ^14,15^.

Although biomarkers have been developed with the aim of diagnosing patients, the focus has shifted to the identification of biomarkers that can determine therapeutic targets, such as theranostic biomarkers ^47,48^, which would allow for more personalized treatment approaches. The 31 FCs discovered in this study are promising candidates for theranostic biomarkers for MDD because they are related to the MDD diagnosis. Future work should investigate whether modulation of FC could be an effective treatment of MDD by using an intervention method with regard to FC, such as functional connectivity neurofeedback training ^47-51^.

## Materials and Methods

### Participants

We used 2 rs-fMRI datasets for the analyses: (1) The “discovery dataset” contained data from 713 participants (564 HCs from 4 sites, 149 MDD patients from 3 sites; Table 1). Each participant underwent a single rs-fMRI session which lasted 10 min. Within the Japanese SRPBS DecNef project, we planned to acquire the rs-fMRI data using a unified imaging protocol (Supplementary Table 1; http://bicr.atr.jp/rs-fmri-protocol-2/). However, there were 2 erroneous phase-encoding directions (P→A and A→P). In addition, different sites had different MRI hardware (Supplementary Table 1). During the rs-fMRI scans, participants were instructed to “Relax. Stay Awake. Fixate on the central crosshair mark, and do not concentrate on specific things”. (2) The “independent validation dataset” contained data from 449 participants (264 HCs from independent 4 sites, 185 MDD patients from independent 4 sites; Table 1). Data were acquired following protocols reported in Supplementary Table 1. The sites used were different from the discovery dataset. Each participant underwent a single rs-fMRI session lasting 5 or 8 min. In both datasets, depression symptoms were evaluated using the BDI-II score obtained from most participants in each dataset. This study was carried out in accordance with the recommendations of the institutional review boards of the principal investigators’ respective institutions (Hiroshima University, Kyoto University, Showa University, University of Tokyo, and Yamaguchi University) with written informed consent from all subjects in accordance with the Declaration of Helsinki. The protocol was approved by the institutional review boards of the principal investigators’ respective institutions (Hiroshima University, Kyoto University, Showa University, University of Tokyo, and Yamaguchi University). Most data utilized in this study can be downloaded publicly from the DecNef Project Brain Data Repository at https://bicr-resource.atr.jp/srpbsopen/ and https://bicr.atr.jp/dcn/en/download/harmonization/. The data availability statements of each site are described in Supplementary Table 1.

### Preprocessing and calculation of the resting state FC matrix

We preprocessed the rs-fMRI data using FMRIPREP version 1.0.8 ^52^. The first 10 s of the data were discarded to allow for T1 equilibration. Preprocessing steps included slice-timing correction, realignment, coregistration, distortion correction using a field map, segmentation of T1-weighted structural images, normalization to Montreal Neurological Institute (MNI) space, and spatial smoothing with an isotropic Gaussian kernel of 6 mm full-width at half-maximum. “Fieldmap-less” distortion correction was performed for the independent validation dataset due to the lack of field map data. For more details on the pipeline, see http://fmriprep.readthedocs.io/en/latest/workflows.html. For 6 participants’ data in the independent validation dataset, the coregistration was unsuccessful, and we therefore excluded these data from further analysis.

#### Parcellation of brain regions

To analyze the data using Human Connectome Project (HCP) style surface-based methods, we used ciftify toolbox version 2.0.2 ^53^. This allowed us to analyze our data, which lacked the T2-weighted image required for HCP pipelines, using an HCP-like surface-based pipeline. Next, we used Glasser’s 379 surface-based parcellations (cortical 360 parcellations + subcortical 19 parcellations) as regions of interest (ROIs), considered reliable brain parcellations ^28^. The BOLD signal time courses were extracted from these 379 ROIs. To facilitate the comparison of our results with previous studies, we identified the anatomical names of important ROIs and the names of intrinsic brain networks that included the ROIs using anatomical automatic labeling (AAL) ^54^ and Neurosynth (http://neurosynth.org/locations/).

#### Physiological noise regression

Physiological noise regressors were extracted by applying CompCor ^55^. Principal components were estimated for the anatomical CompCor (aCompCor). A mask to exclude signals with a cortical origin was obtained by eroding the brain mask and ensuring that it contained subcortical structures only. Five aCompCor components were calculated within the intersection of the subcortical mask and union of the CSF and WM masks calculated in the T1-weighted image space after their projection to the native space of functional images in each session. To remove several sources of spurious variance, we used a linear regression with 12 regression parameters, such as 6 motion parameters, average signals over the whole brain, and 5 aCompCor components.

#### Temporal filtering

A temporal bandpass filter was applied to the time series using a first-order Butterworth filter with a pass band between 0.01 Hz and 0.08 Hz to restrict the analysis to low-frequency fluctuations, which are characteristic of rs-fMRI BOLD activity ^56^.

#### Head motion

Framewise displacement (FD) ^57^ was calculated for each functional session using Nipype (https://nipype.readthedocs.io/en/latest/). FD was used in the subsequent scrubbing procedure. To reduce spurious changes in FC from head motion, we removed volumes with FD > 0.5 mm, as proposed in a previous study ^57^. The FD represents head motion between 2 consecutive volumes as a scalar quantity (i.e., the summation of absolute displacements in translation and rotation). Using the aforementioned threshold, 6.3% ± 13.5 volumes (mean ± SD) were removed per rs-fMRI session in all datasets. If the ratio of the excluded volumes after scrubbing exceeded the mean + 3 SD, participants were excluded from the analysis. As a result, 32 participants were removed from all datasets. Thus, we included 683 participants (545 HCs, 138 MDD patients) in the discovery dataset and 440 participants (259 HCs, 181 MDD patients) in the independent validation dataset for further analysis.

#### Calculation of FC matrix

FC was calculated as the temporal correlation of rs-fMRI BOLD signals across 379 ROIs for each participant. There are a number of different candidates to measure FC, such as the tangent method and partial correlation; however, we used a Pearson’s correlation coefficient because they are the most commonly used values in previous studies. Fisher’s z-transformed Pearson’s correlation coefficients were calculated between the preprocessed BOLD signal time courses of each possible pair of ROIs and used to construct 379 × 379 symmetrical connectivity matrices in which each element represents a connection strength between 2 ROIs. We used 71,631 FC values [(379 × 378)/2] of the lower triangular matrix of the connectivity matrix for further analysis.

#### Control of site differences

Next, we used a traveling subject harmonization method to control for site differences in FC in the discovery dataset. This method enabled us to subtract pure site differences (measurement bias) which are estimated from the traveling subject dataset wherein multiple participants travel to multiple sites to assess measurement bias. The participant factor (***p***), measurement bias (***m***), sampling biases (***s***_hc_, ***s***_mdd_), and psychiatric disorder factor (***d***) were estimated by fitting the regression model to the FC values of all participants from the discovery dataset and the traveling subject dataset. For each connectivity, the regression model can be written as follows:

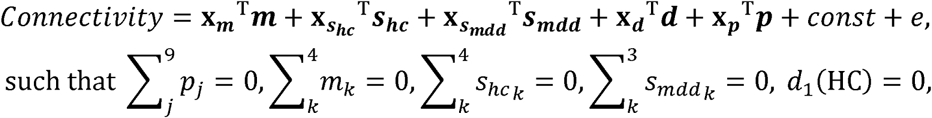

in which ***m*** represents the measurement bias (4 sites × 1), ***s***_***hc***_ represents the sampling bias of HCs (4 sites × 1), ***s***_***mdd***_ represents the sampling bias of patients with MDD (3 sites × 1), ***d*** represents the disorder factor (2 × 1), ***p*** represents the participant factor (9 traveling subjects × 1), *const* represents the average functional connectivity value across all participants from all sites, and *e*∼ 𝒩 (0, *γ*^−1^) represents noise. Measurement biases were removed by subtracting the estimated measurement biases. Thus, the harmonized functional connectivity values were set as follows:

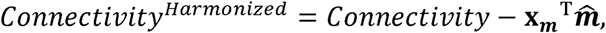

in which 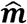 represents the estimated measurement bias. More detailed information has been previously described ^26^.

We used the ComBat harmonization method ^29-32^ to control for site differences in FC in the independent validation dataset because we did not have a traveling subject dataset for those sites. We performed harmonization to correct only for the site difference using information on MDD diagnosis, BDI score, age, sex, and dominant hand as auxiliary variables in ComBat. Notably, compared with the conventional regression method, the ComBat method is a more advanced method to control for site effects ^29-32^.

### Constructing the MDD classifier using the discovery dataset

We constructed a brain network marker for MDD that distinguished between HCs and MDD patients using the discovery dataset based on 71,631 FC values. To construct the network marker, we applied a machine learning technique. Although SVM is often used as a classifier, SVM is not suitable for investigating the contribution of explanatory variables because it is difficult to calculate the contribution of each explanatory variable. Based on our previous study ^13^, we assumed that psychiatric disorder factors were not associated with whole brain connectivity, but rather with a specific subset of connections. Therefore, we conducted logistic regression analyses using the LASSO method to select the optimal subset of FCs ^34^. A logistic function was used to define the probability of a participant belonging to the MDD class as follows:

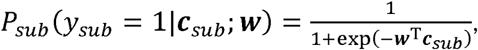

in which ***y***_***sub***_ represents the class label (MDD, *y* = 1; HC, *y* = 0) of a participant, ***c***_***sub***_ represents an FC vector for a given participant, and *w* represents the weight vector. The weight vector *w* was determined to minimize

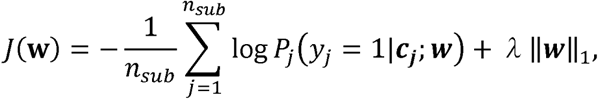

in which 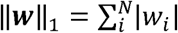 and λ represent hyperparameters that control the amount of shrinkage applied to the estimates. To estimate weights of the logistic regression and a hyperparameter λ, we conducted a nested cross validation procedure (Fig. 2). In this procedure, we first divided the whole discovery dataset into a training set (9 folds of 10 folds) which used for training a model and a test set (a fold of 10 folds) for testing the model. To minimize bias due to the differences in the numbers of MDD patients and HCs, we used an undersampling method ^35^. Almost 125 MDD patients and 125 HCs were randomly sampled from the training set, and classifier performance was tested using the test set. Since only a subset of training data is used after undersampling, we repeated the random sampling procedure 10 times (i.e., subsampling). We then fitted a model to each subsample while tuning a regularization parameter in the inner loop of the nested cross validation, resulting in 10 classifiers. For the inner loop, we used the “*lassoglm*” function in MATLAB (R2016b, Mathworks, USA) and set “NumLambda” to 25 and “CV” to 10. In this inner loop, we first calculated a value of λ just large enough such that the only optimal solution is the all-zeroes vector. A total of 25 values of λ were prepared at equal intervals from 0 to λ_max_ and the λ was determined according to the one-standard-error-rule in which we selected the largest λ within the standard deviation of the minimum prediction error (among all λ) ^27^. The mean classifier output value (diagnostic probability) was considered indicative of the classifier output. Diagnostic probability values > 0.5 were considered indicative of MDD patients. We calculated the AUC using the “*perfcurve*” function in MATLAB. In addition, we calculated the accuracy, sensitivity, specificity, PPV, and NPV. Furthermore, we evaluated classifier performance for the unbalanced dataset using the MCC ^36,37^, which takes into account the ratio of the confusion matrix size.

### BDI score regression model in the discovery dataset

We constructed a linear regression model to predict the BDI score using the discovery dataset based on 71,631 FC values. To construct the linear regression model, we applied a machine-learning technique to participants with BDI scores in the discovery dataset. Although SVR is often used as a regression model, SVR is not suitable for investigating the contribution of explanatory variables because it is difficult to calculate the contribution of each explanatory variable. Therefore, we employed linear regression using the LASSO method as follows:

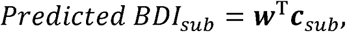

in which *Predicted BDI*_*sub*_ represents the BDI score of a participant; ***c***_*sub*_ represents an FC vector for a given participant, and ***w*** represents the weight vector of the linear regression. The prediction model was constructed while feature selection using the embedded method with LASSO was performed (Fig. 2) ^39^. We conducted a 10-fold CV procedure for this regression model. We constructed a regression model using the combination of FC values selected in all 10 folds in the training dataset (Fig. 2). This caused information leakage across the folds; therefore, the training dataset may be overfitting. This issue meant that it was important to confirm generalization performance by applying this regression model to an independent validation dataset, as described below. Finally, we calculated the mean absolute error (MAE) and Pearson’s correlation coefficients between the predicted and measured BDI scores.

### Generalization performance of the classifier and regression model

We tested the generalizability of the classifier and regression model using an independent validation dataset. We created 100 classifiers of MDD (10-fold × 10 subsamples); therefore, we applied all trained classifiers to the independent validation dataset. Next, we averaged the 100 outputs (diagnostic probability) for each participant and considered the participant as a patient with MDD if the averaged diagnostic probability value was >0.5. In contrast, we created the BDI regression model using all the discovery dataset samples; therefore, we applied the trained regression model to the independent validation dataset and considered its output as the predicted BDI score.

To test the statistical significance of the MDD classifier performance, we performed a permutation test. We permuted the diagnostic labels of the discovery dataset and conducted a 10-fold CV and 10-subsampling procedure. Next, we took an average of the 100 outputs (diagnostic probability); a mean diagnostic probability value > 0.5 was considered indicative of a diagnosis of MDD. We repeated this permutation procedure 100 times and calculated the AUC and MCC as the performance of each permutation. We also performed a permutation test for the BDI regression model. We permuted the BDI scores of the discovery dataset, conducted a 10-fold CV, and repeated this permutation procedure 100 times.

### Identification of the FCs linked to diagnosis and symptoms

We examined resting-state functional connectivity for MDD diagnosis and depression symptoms by extracting the important FCs related to the MDD classifier and BDI regression model, respectively. Briefly, we counted the number of times an FC was selected by LASSO during the 10-fold CV. We considered that this FC was important if this number was significantly higher than chance, according to a permutation test. We permuted the diagnostic labels of the discovery dataset and conducted a 10-fold CV and 10-subsampling procedure. We then used the number of counts for each connection selected by the sparse algorithm during 10 subsampling x 10 CV(max 100 times) as a statistic in every permutation dataset. To control the multiple comparison problem, we set a null distribution as the max distribution of the number of counts over all functional connections and set our statistical significance to a threshold (*p* < 0.05, one-sided). FCs selected ≥ 17 times of 100 times were regarded as diagnostically important. We also performed a permutation test for the BDI regression model. We permuted the BDI scores of the discovery dataset, conducted a 10-fold CV, and repeated this permutation procedure 100 times. FCs selected ≥ 1 times of 10 times were regarded as relevant to depression symptoms.

## Acknowledgements

We would like to thank Editage (www.editage.com) for English language editing.

## Funding

This study was conducted under the “Development of BMI Technologies for Clinical Application” of the Strategic Research Program for Brain Sciences, and the contract research Grant Number JP18dm0307008, supported by the Japan Agency for Medical Research and Development (AMED). This study was also partially supported by the ImPACT Program of the Council for Science, Technology and Innovation (Cabinet Office, Government of Japan). This study was also supported by AMED under Grant Number JP18dm0307001 & JP18dm0307004, JSPS KAKENHI 16H06280 (Advanced Bioimaging Support), and the International Research Center for Neurointelligence (WPI-IRCN) at The University of Tokyo Institutes for Advanced Study (UTIAS). H.I. was partially supported by JSPS KAKENHI 26120002, 18H01098, and 18H05302.

## Author contributions

A.Y. and M.K. designed the study. T.Y., N.Y., A.K., N.O., T.I., R.H., H.M., N.I., M.T., G.O., H.Y., K.H., K.M., S.T., K.K., N.K., H.T., and Y.O. recruited participants for the study, collected their clinical and imaging data, and constructed the database. A.Y. performed data preprocessing. A.Y. and O.Y. performed analysis under the supervision of M.K. and H.I.; A.Y., M.K., O.Y., and H.I. primarily wrote the manuscript.

## Competing financial interests

Competing financial interests: M.K., N.Y., R.H., H.I., N.K. and K.K are inventors of a patent owned by Advanced Telecommunications Research (ATR) Institute International related to the present work [PCT/JP2014/061543 (WO2014178322)]. M.K., N.Y., R.H., N.K. and K.K. are inventors of a patent owned by ATR Institute International related to the present work [PCT/JP2014/061544 (WO2014178323)]. M.K. and N.Y. are inventors of a patent application submitted by ATR Institute International related to the present work [JP2015-228970]. A.Y. and M.K. are inventors of a patent application submitted by ATR Institute International related to the present work [JP2018-192842].

## Supplementary Materials

### Supplementary Note 1

#### Prediction performance using SVM

Since the performance levels of the prediction models were acceptable, we further tried a support vector machine (SVM) for classification. However, the performance was not improved compared to that in LASSO. In the discovery dataset, the classifier of SVM separated major depressive disorder (MDD) and healthy control (HC) populations with an accuracy of 71%. The corresponding AUC sensitivity and specificity were 0.78, 72% and 70%, respectively. This approach led to a Matthews correlation coefficient (MCC) of 0.35. In the independent validation dataset, the classifier of SVM separated the MDD and HC populations with an accuracy of 70%. The corresponding AUC, sensitivity, and specificity were 0.75, 62%, and 76%, respectively. This approach led to an MCC of 0.38.

### Supplementary Note 2

#### Differences in prediction performance among imaging sites

To investigate whether the prediction performances were different among imaging sites in the independent validation dataset, we calculated the 95% confidence interval (CI) of discrimination performances (area under the curve [AUC], accuracy, sensitivity, and specificity) in every imaging site using a bootstrap method. We repeated the bootstrap procedure 1,000 times and calculated the 95% CI for every site. We then checked whether there is a site whose CI does not overlap with the CIs of other imaging sites. We could not find such an imaging site, suggesting no significant systematic difference. However, we noted that the sensitivity at the HUH site was inferior to that at the two other imaging sites (Supplementary Fig. 1: CI of sensitivity in the HUH does not overlap with CI in the HRC or UYA). We discussed the differences in performance among imaging sites in the Discussion section in the main texts.

**Supplementary Figure 1:**
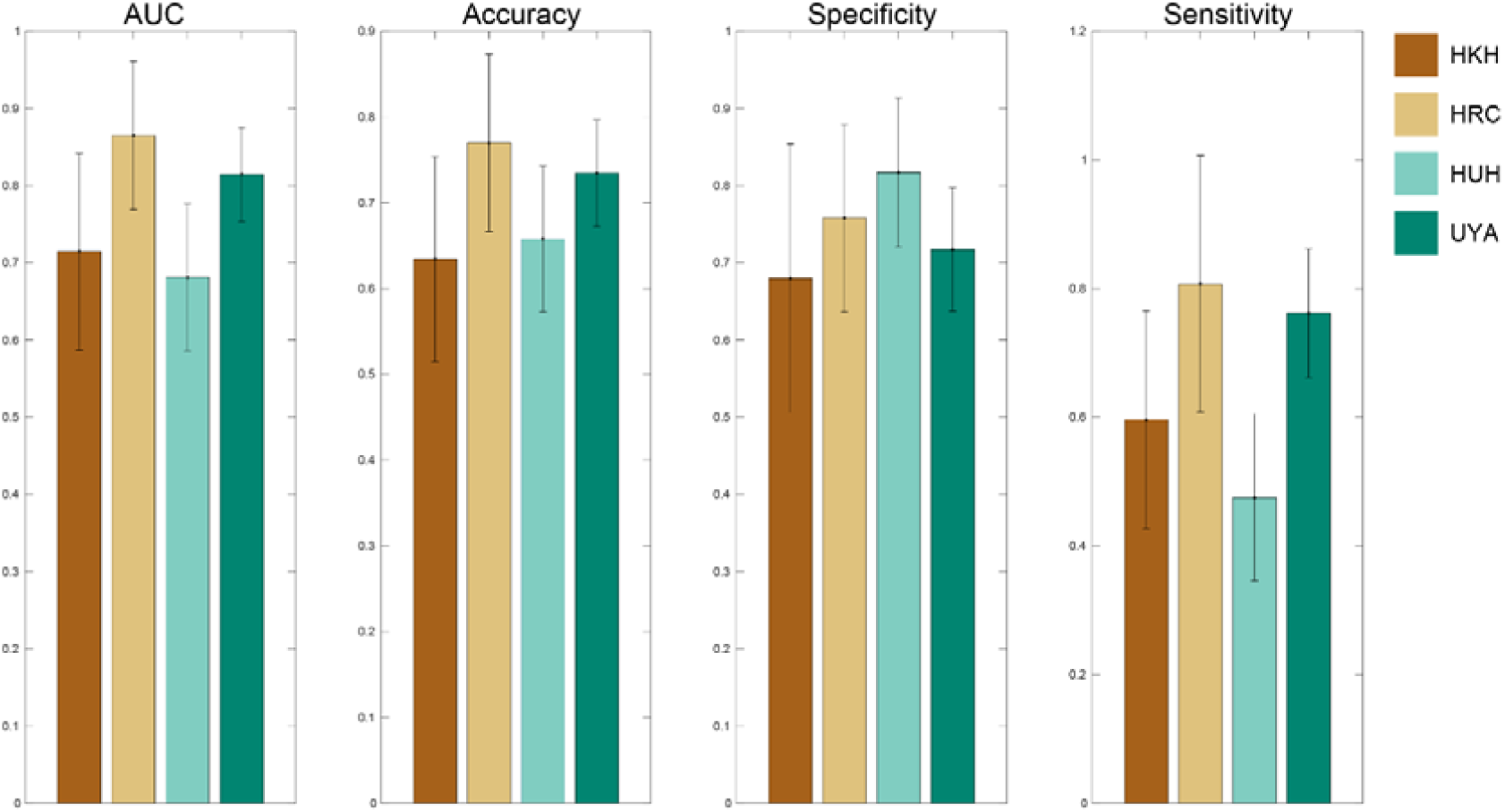
Bootstrap prediction performances in the independent validation dataset. Prediction performances of the major depressive disorder (MDD) classifier in the independent validation dataset in each site. Each color bar indicates each site. Error bar shows the 95 % confidence interval from the bootstrap. AUC: Area under the curve, HKH: Hiroshima Kajikawa Hospital; HRC: Hiroshima Rehabilitation Center; HUH: Hiroshima University Hospital; UYA: Yamaguchi University.

### Supplementary Note 3

#### Generalization of the classifiers to other disorders

In addition to a brain network marker of MDD, we developed brain network markers of schizophrenia (SCZ) and autism spectrum disorder (ASD) using the same method as in the main text. We sought to investigate and confirm the spectral structure among the disorders as revealed by previous studies (*12-16* in the main script). For example, if the MDD classifier predicts patients with a different disorder as MDD patients, then the probability of diagnosis for patients with that disorder should be over 0.5. In this case, we may say that the patients possess some degree of MDD-ness and that this disorder is related to MDD according to the imaging biological dimension.

Specifically, we first developed SCZ and ASD markers that distinguished between HCs and patients. We used 564 HCs from the discovery dataset in the main text, 102 SCZ patients from 3 sites, and 121 ASD patients from 2 sites (Supplementary Table 5). Data were acquired using the same protocols as for the discovery dataset. We tested the generalizability of the SCZ marker using an independent validation dataset for SCZ patients (52 SCZ patients and 75 HCs from one site, Supplementary Table 5). Since we did not have an independent validation dataset for ASD patients, we tested the performance of the ASD marker using the 10-fold CV. We achieved acceptable performance for both the SCZ marker (Discovery dataset: AUC = 0.85, accuracy = 78%, sensitivity = 75%, specificity = 79%, Independent validation dataset: AUC = 0.89, accuracy = 80%, sensitivity = 81%, specificity = 80%) and ASD marker (Discovery dataset: AUC = 0.76, accuracy = 65%, sensitivity = 70%, specificity = 64%). We then applied these brain network markers to other disorder patients. We computed the probability of diagnosis in the MDD classifier, that is, the MDD-ness of individual patient within the SCZ and ASD data, and vice versa (Supplementary Figure 2).

As a result, we found that SCZ patients have high MDD-ness (accuracy = 76%, *p* = 2.0*10^−12^, two-way binomial test) and ASD-ness (accuracy = 68%, *p* = 2.1*10^−4^, two-way binomial test). On the other hand, MDD patients did not have high SCZ-ness (accuracy = 46%, *p* = 0.35, two-way binomial test) or ASD-ness (accuracy = 57%, *p* = 0.11, two-way binomial test), and ASD patients did not have high SCZ-ness (accuracy = 42%, *p* = 0.10, two-way binomial test) or MDD-ness (accuracy = 55%, *p* = 0.20, two-way binomial test).

**Supplementary Figure 2:**
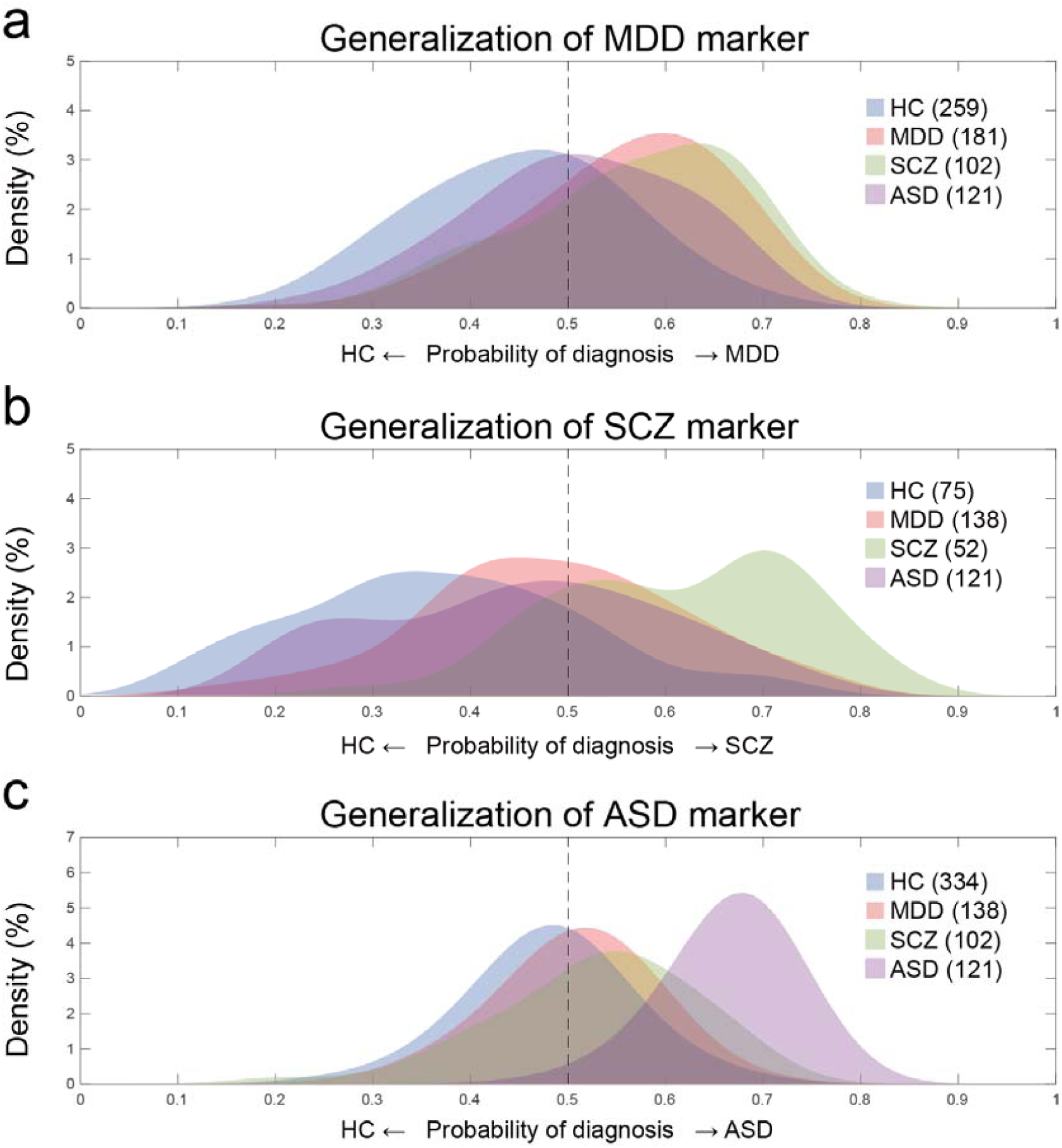
Generalization of the classifiers to other psychiatric disorders. Density distributions of the probability of diagnosis obtained by applying (a) the MDD marker, (b) SCZ marker, and (c) ASD marker to the HC, MDD, SCZ, and ASD patients. In each panel, the patient distribution and the healthy control distribution are plotted separately, with the colored areas representing one or the other. The numbers in parentheses next to HC, MDD, ASD, and SCZ in each panel indicate the number of subjects in the distributions. The independent validation dataset was used in (a) and (b). Healthy controls in (a), (b), and (c) were scanned at the same sites as their corresponding patient data. HC: healthy control; MDD: major depressive disorder; ASD: autism spectrum disorder; SCZ: schizophrenia.

**Supplementary Table 1.**
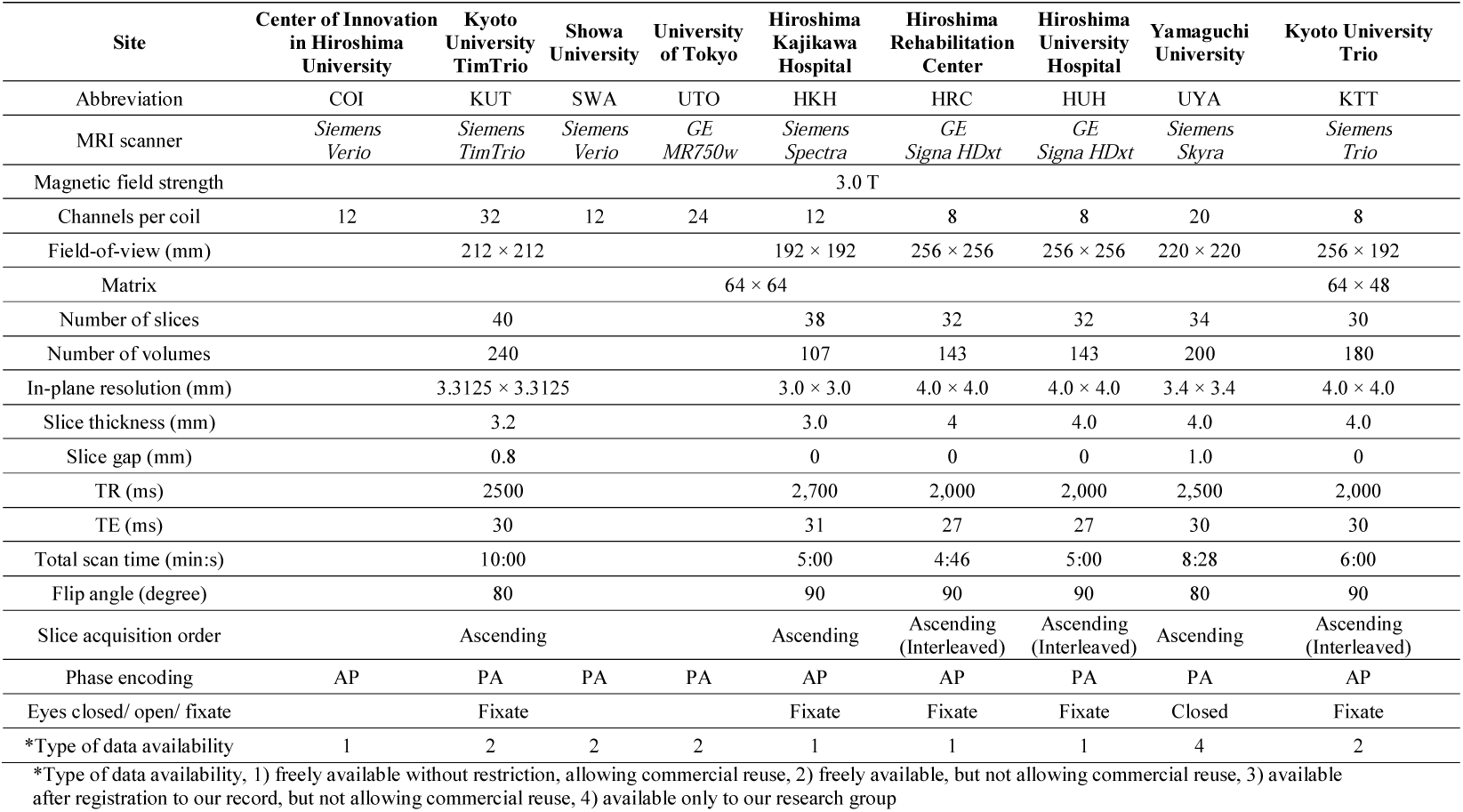
Imaging protocols for resting-state fMRI in both.

**Supplementary Table 2.**
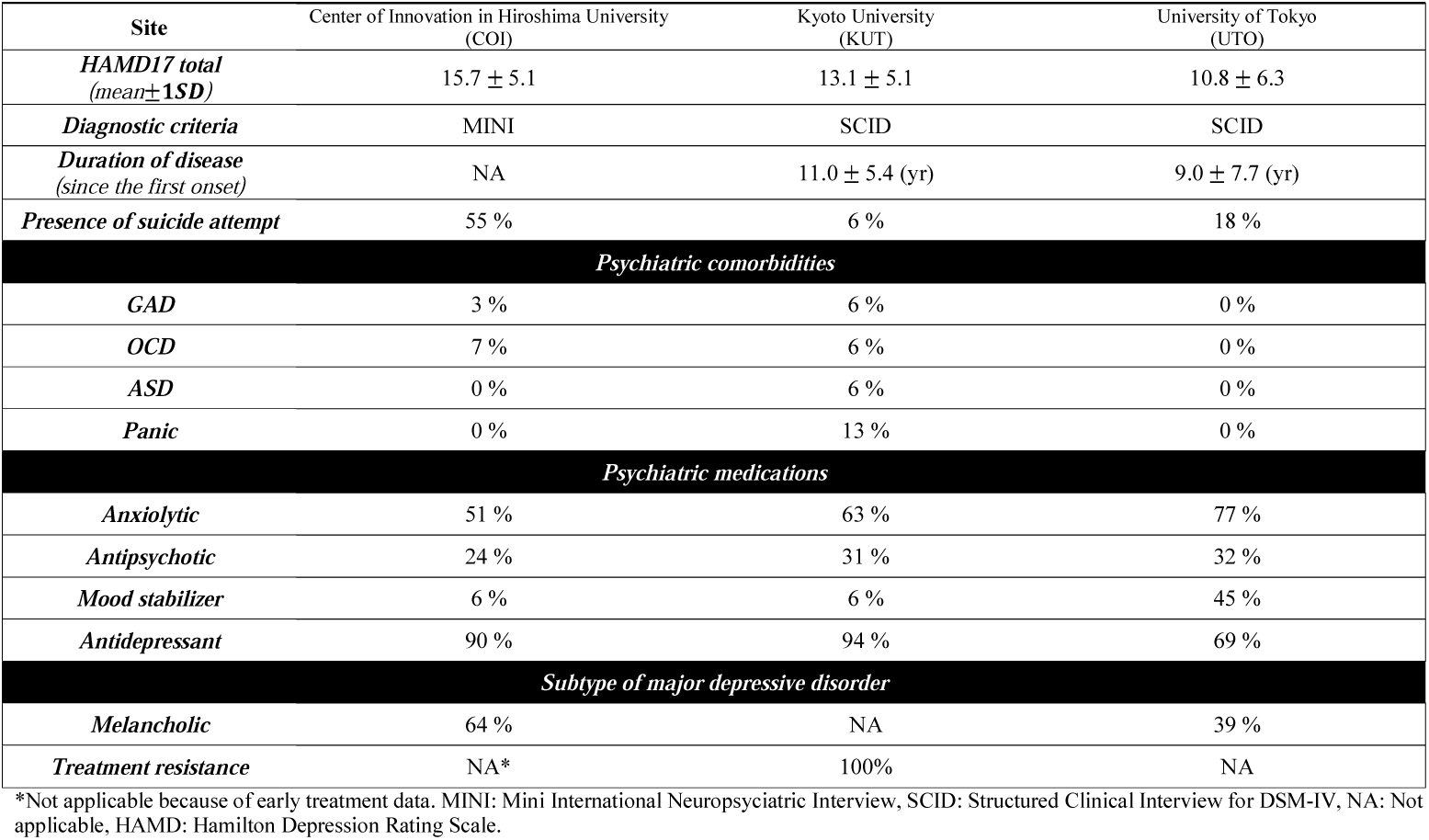
Clinical characteristics of major depressive disorder patients in the discovery dataset.

|  |  |  |  |
| --- | --- | --- | --- |
| Site | Center of Innovation in Hiroshima University (COI) | Kyoto University (KUT) | University of Tokyo (UTO) |
| *HAMD17 total (mean±1SD)* | 15.7 ± 5.1 | 13.1 ± 5.1 | 10.8 ± 6.3 |
| *Diagnostic criteria* | MINI | SCID | SCID |
| *Duration of disease (since the first onset)* | NA | 11.0 ± 5.4 (yr) | 9.0 ± 7.7 (yr) |
| *Presence of suicide attempt* | 55 % | 6 % | 18 % |
| *Psychiatric comorbidities* |  |  |  |
| *GAD* | 3 % | 6 % | 0 % |
| *OCD* | 7 % | 6 % | 0 % |
| *ASD* | 0 % | 6 % | 0 % |
| *Panic* | 0 % | 13 % | 0 % |
| *Psychiatric medications* |  |  |  |
| *Anxiolytic* | 51 % | 63 % | 77 % |
| *Antipsychotic* | 24 % | 31 % | 32 % |
| *Mood stabilizer* | 6 % | 6 % | 45 % |
| *Antidepressant* | 90 % | 94 % | 69 % |
| *Subtype of major depressive disorder* |  |  |  |
| *Melancholic* | 64 % | NA | 39 % |
| *Treatment resistance* | NA* | 100% | NA |

\*Not applicable because of early treatment data. MINI: Mini International Neuropsychiatric Interview, SCID: Structured Clinical Interview for DSM-IV, NA: Not applicable, HAMD: Hamilton Depression Rating Scale.

**Supplementary Table 3.**
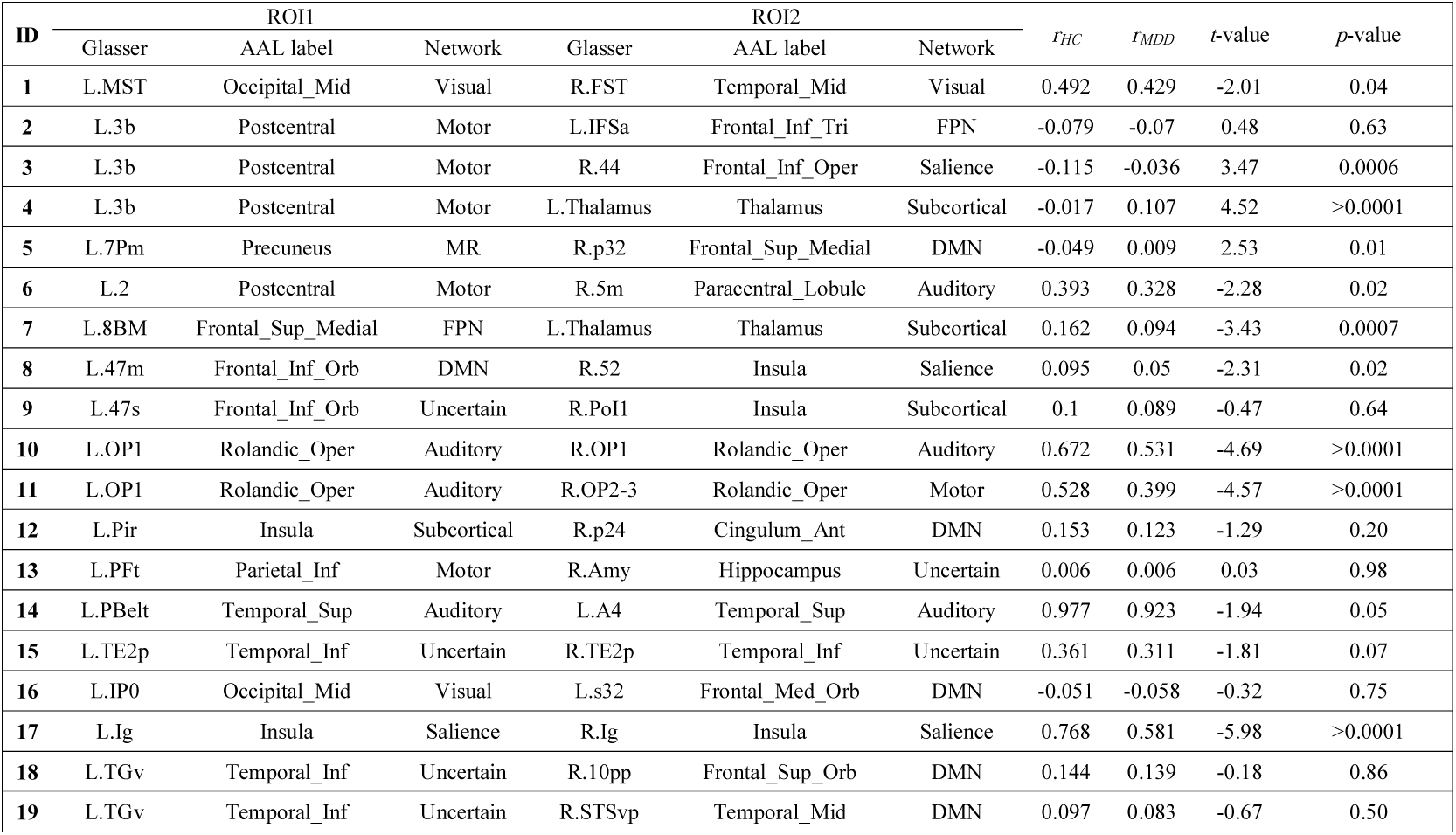
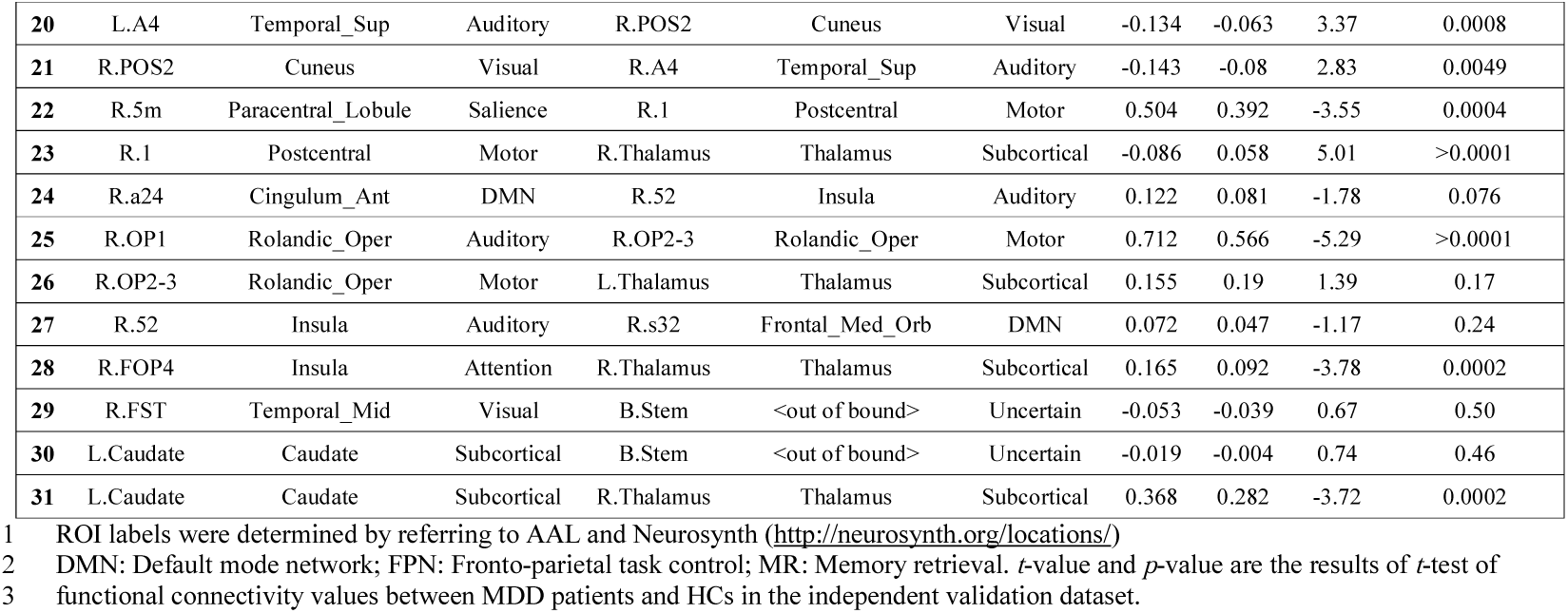
Description of important FCs.

**Supplementary Table 4.** All functional connections related to only BDI score regression model.

|  |  |  |  |  |  |  |  |  |
| --- | --- | --- | --- | --- | --- | --- | --- | --- |
| ID | ROI1 |  |  | ROI2 |  |  | *r*-value<br>with BDI<br>(Discovery) | *r*-value<br>with BDI<br>(Validation) |
|  | Glasser | AAL label | Network | Glasser | AAL label | Network |  |  |
| 1 | L.3b | Postcentral_L | Motor | R.Thalamus | Thalamus_R | Subcortical | 0.28 | 0.04 |
| 2 | L.POS1 | Precuneus_L | DMN | R.HC | Hippocampus_R | Uncertain | 0.22 | -0.04 |
| 3 | L.9m | Frontal_Sup_Medial_L | DMN | R.9m | Frontal_Sup_Medial_R | DMN | -0.21 | -0.06 |
| 4 | L.AAIC | Insula_L | Uncertain | R.p24 | Cingulum_Ant_R | DMN | -0.21 | -0.02 |
| 5 | L.TGd | Temporal_Pole_Mid_L | DMN | R.STSvp | Temporal_Mid_R | DMN | -0.21 | -0.08 |
| 6 | L.FST | Temporal_Mid_L | Attention | R.RSC | Cingulum_Post_R | DMN | 0.21 | 0.04 |
| 7 | L.VMV2 | Lingual_L | DMN | R.VMV2 | Lingual_R | Visual | -0.23 | 0.02 |
| 8 | L.FOP5 | Insula_L | Salience | R.FFC | Fusiform_R | Uncertain | 0.18 | -0.01 |
| 9 | L.PI | Temporal_Sup_L | Attention | R.TE1m | Temporal_Mid_R | Uncertain | -0.20 | -0.01 |
| 10 | R.a24 | Cingulum_Ant | DMN | R.52 | Insula | Auditory | -0.24 | 0.06 |
| 11 | R.52 | Insula | Auditory | R.s32 | Frontal_Med_Orb | DMN | -0.24 | 0.03 |
| 12 | R.AIP | Parietal_Inf_R | FPN | R.LBelt | Temporal_Sup_R | Auditory | 0.18 | -0.05 |
| 13 | L.Putamen | Putamen_L | Subcortical | R.Putamen | Putamen_R | Subcortical | -0.25 | 0.03 |

ROI labels were determined by referring to AAL and Neurosynth (http://neurosynth.org/locations/)
DMN: Default mode network; FPN: Fronto-parietal task control; MR: Memory retrieval.

**Supplementary Table 5.**
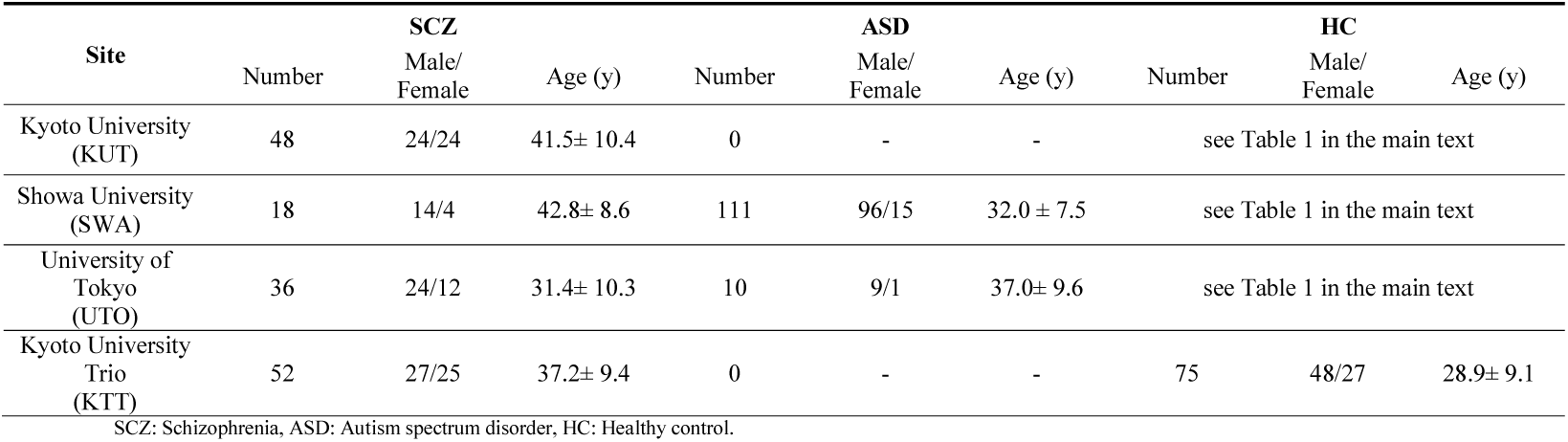
Demographic characteristics of participants in both datasets.

**Supplementary Table 6.** Imaging protocols for resting state fMRI in the traveling subject dataset.

| Site | ATR<br>TimTrio | ATR<br>Verio | Center of<br>Innovation in<br>Hiroshima<br>University | Hiroshima<br>University<br>Hospital | Hiroshima<br>Kajikawa<br>Hospital | Kyoto<br>Prefectural<br>University of<br>Medicine | Showa<br>University | Kyoto<br>University<br>TimTrio | Kyoto<br>University<br>Skyra | University<br>of Tokyo | Yaesu-clinic<br>scanner 1 | Yaesu-clinic<br>scanner 2 |
| --- | --- | --- | --- | --- | --- | --- | --- | --- | --- | --- | --- | --- |
| Abbreviation | ATT | ATV | COI | HUH | HKH | KPM | SWA | KUT | KUS | UTO | YC1 | YC2 |
| MRI scanner | *Siemens*<br>*TimTrio* | *Siemens*<br>*Verio* | *Siemens*<br>*Verio* | *GE*<br>*Signa HDxt* | *Siemens*<br>*Spectra* | *Philips*<br>*Achieva* | *Siemens*<br>*Verio* | *Siemens*<br>*TimTrio* | *Siemens*<br>*Skyra* | *GE*<br>*MR750W* | *Philips*<br>*Achieva* | *Philips*<br>*Achieva* |
| The number of scans | 132 | 27 | 27 | 18 | 18 | 27 | 27 | 27 | 27 | 27 | 27 | 27 |
| Magnetic field strength |  |  |  |  |  | 3T |  |  |  |  |  |  |
| Number of channels per coil | 12 | 12 | 12 | 8 | 12 | 8 | 12 | 32 | 32 | 24 | 8 | 8 |
| Field-of-view (mm) |  |  |  |  |  | 212 x 212 |  |  |  |  |  |  |
| Matrix |  |  |  |  |  | 64 × 64 |  |  |  |  |  |  |
| Number of slices | 40 | 39 | 40 | 35 | 35 | 40 | 40 | 40 | 40 | 40 | 40 | 40 |
| Number of volumes |  |  |  |  |  | 240 |  |  |  |  |  |  |
| In-plane resolution (mm) |  |  |  |  |  | 3.3125 × 3.3125 |  |  |  |  |  |  |
| Slice thickness (mm) |  |  |  |  |  | 3.2 |  |  |  |  |  |  |
| Slice gap (mm) |  |  |  |  |  | 0.8 |  |  |  |  |  |  |
| TR (ms) |  |  |  |  |  | 2,500 |  |  |  |  |  |  |
| TE (ms) |  |  |  |  |  | 30 |  |  |  |  |  |  |
| Total scan time (min:s) |  |  |  |  |  | 10:00 |  |  |  |  |  |  |
| Flip angle (deg) |  |  |  |  |  | 80 |  |  |  |  |  |  |
| Slice acquisition order |  |  |  |  |  | Ascending |  |  |  |  |  |  |
| Phase encoding | PA | PA | AP | PA | PA | AP | PA | PA | AP | PA | AP | AP |
| Eye closed / fixate |  |  |  |  |  | Fixate |  |  |  |  |  |  |
| Field map | ✓ | ✓ | ✓ | - | - | ✓ | ✓ | ✓ | ✓ | ✓ | ✓ | - |

1 Nine healthy participants (all male participants; age range, 24–32 years; mean age, $27 \pm 2.6$ years) were scanned at each of 12 sites, producing a total of 411 scan 2 sessions. Each participant underwent three rs-fMRI sessions of 10 min each at nine sites, two sessions of 10 min each at two sites (HKH & HUH), and five cycles 3 (morning, afternoon, next day, next week, next month) consisting of three 10-minute sessions each at a single site (ATT). In the latter situation, one participant 4 underwent four rather than five sessions at the ATT site due to poor physical condition. Thus, a total of 411 sessions were conducted. UTO: University of Tokyo; HUH: Hiroshima University Hospital; KUT: Siemens TimTrio scanner at Kyoto University; ATT: Siemens TimTrio scanner at Advanced Telecommunications Research Institute International; ATV: Siemens Verio scanner at Advanced Telecommunications Research Institute International; SWA: Showa University; HKH: Hiroshima Kajikawa Hospital; COI: Center of Innovation in Hiroshima University; KUS: Siemens Skyra scanner at Kyoto University; KPM: Kyoto Prefectural University of Medicine; YC1: Yaesu Clinic 1; YC2: Yaesu Clinic 2; TR: repetition time; TE: echo time.

